# CXIP4 depletion causes early lethality and pre-mRNA missplicing in Arabidopsis

**DOI:** 10.1101/2024.06.06.597795

**Authors:** Uri Israel Aceituno-Valenzuela, Sara Fontcuberta-Cervera, Rosa Micol-Ponce, Raquel Sarmiento-Mañús, Alejandro Ruiz-Bayón, María Rosa Ponce

## Abstract

Zinc knuckle (ZCCHC) motif-containing proteins are present in unicellular and multicellular eukaryotes and most ZCCHC proteins with known functions participate in the metabolism of various classes of RNA, such as mRNAs, ribosomal RNAs, and microRNAs. The Arabidopsis (*Arabidopsis thaliana*) genome encodes 69 ZCCHC-containing proteins, but the functions of most remain unclear. One of these proteins is CAX-INTERACTING PROTEIN 4 (CXIP4), which has been classified as a PTHR31437 family member, along with human SREK1-interacting protein 1 (SREK1IP1), which is thought to function in pre-mRNA splicing and RNA methylation. Metazoan SREK1IP1-like and plant CXIP4-like proteins only share a ZCCHC motif, and their functions remain almost entirely unknown. We studied two loss-of-function alleles of Arabidopsis *CXIP4*, the first mutations in PTHR31437 family genes described to date: *cxip4-1* is likely null and shows early lethality, and *cxip4-2* is hypomorphic and viable, with pleiotropic morphological defects. The *cxip4-2* mutant exhibited deregulation of defense genes and upregulation of transcription factor encoding genes, some of which might explain its developmental defects. This mutant also exhibited increased intron retention events, and the specific functions of misspliced genes, such as those involved in “gene silencing by DNA methylation” and “mRNA polyadenylation factor” suggest that CXIP4 has additional functions. The CXIP4 protein localizes to the nucleus in a pattern resembling nuclear speckles, which are rich in splicing factors. Therefore, *CXIP4* is required for plant survival and proper development, and mRNA maturation.

## INTRODUCTION

Zinc knuckles are 18-residue CCHC-type zinc finger (ZCCHC) motifs with the conserved sequence CX_2_CX_4_HX_4_C. These motifs are present in proteins of unicellular and multicellular eukaryotes, from yeast to plants and animals, but are not found in bacteria. Most ZCCHC-containing proteins with known functions are involved in the metabolism of different classes of RNAs, including messenger RNAs (mRNAs), ribosomal RNAs (rRNAs), and microRNAs (miRNAs). The Arabidopsis (*Arabidopsis thaliana*) genome encodes 69 ZCCHC-containing proteins, only a few of which have been functionally characterized at some level (reviewed in Aceituno-Valenzuela et al., 2020).

The AT2G28910 Arabidopsis gene encodes the ZCCHC-containing protein CAX-INTERACTING PROTEIN 4 (CXIP4). Arabidopsis CXIP4 activated the H^+^/Ca^2+^ antiporter CAX1 when expressed in yeast (Cheng et al., 2004). CXIP4 has been classified as a member of the Splicing regulatory glutamine/lysine-rich protein 1 (SREK1)-interacting protein 1 (SREK1IP1) family, which has rarely been studied. Human SREK1IP1 (also known as P18SRP or SFRS12IP1) interacts with the serine-arginine (SR)-rich splicing regulatory protein SREK1, as revealed in yeast two-hybrid (Y2H) assays. In turn, SREK1 interacts with other SR proteins, modulating splice site (SS) selection during alternative splicing (AS) events (Heese et al., 2004). Downregulation of human *SREK1IP1* by RNA interference promoted cell proliferation, migration, and invasion in tumor cell lines (Akiyama et al., 2016), pointing to its role in suppressing tumor growth. Human SREK1IP1 has also been identified as an interactor of METHYLTRANSFERASE LIKE 16 (METTL16; Covelo-Molares et al., 2021), one of the two enzymes that catalyze the deposition of N^6^-methyladenosine (m^6^A) epitranscriptomic marks onto mRNAs and U6 small nuclear RNA (snRNA) (Warda et al., 2017). Dominant frameshift alleles of *SREK1IP1* have been associated with congenital anosmia (Kamarck et al., 2024).

We previously identified Arabidopsis CXIP4 in a Y2H-based screen for interactors of MORPHOLOGY OF ARGONAUTE1-52 SUPPRESSED2 (MAS2; Sánchez-García et al., 2015), which is the putative Arabidopsis ortholog of human NF-kappa-B-activating protein (NKAP; Chen et al., 2003). The multifunctional protein NKAP was recently identified as an exon ligation factor in the post-catalytic (C*) spliceosome (Fica et al.,2019), and as an m^6^A epitranscriptomic mark reader during miRNA and mRNA maturation (Zhang et al., 2019; Sun et al., 2022).

Overexpression of Arabidopsis *CXIP4* conferred resistance to dithiothreitol (DTT) and tunicamycin (TM), two drugs that induce endoplasmic reticulum stress and trigger the unfolded protein response (UPR; Hossain et al., 2016). One of the four wheat (*Triticum aestivum*) CXIP4 co-orthologs functions in calcium-mediated plant immune responses during Fusarium infection (Chen et al., 2022). Despite these findings, the precise functions of plant CXIP4-like and metazoan SREK1IP1-like proteins remain elusive, and loss-of-function mutants have not been characterized in any eukaryote.

Here, we performed functional analysis of Arabidopsis CXIP4 based on two T-DNA insertional alleles, *cxip4-1* and *cxip4-2*. Our findings provide molecular and genetic evidence for the essential role of CXIP4 in Arabidopsis development and survival, as well as its involvement in pre-mRNA splicing.

## RESULTS

### *CXIP4* is an essential gene, required for plant survival and proper development

Most ZCCHC-containing proteins studied to date participate in RNA metabolism (Aceituno-Valenzuela et al., 2020). Arabidopsis *CXIP4* (AT2G28910) is a single-copy gene with two exons (Fig.1A), the second of which encodes a protein of 332 amino acids (aa) containing a ZCCHC motif (residues 81 to 98; Supplementary Fig. S1). CXIP4 also contains a region rich in arginine (R; residues 198 to 289; Supplementary Fig. S1), annotated as an R-rich region in the UniProtKB database (https://www.uniprot.org/uniprotkb).

**Figure 1.**
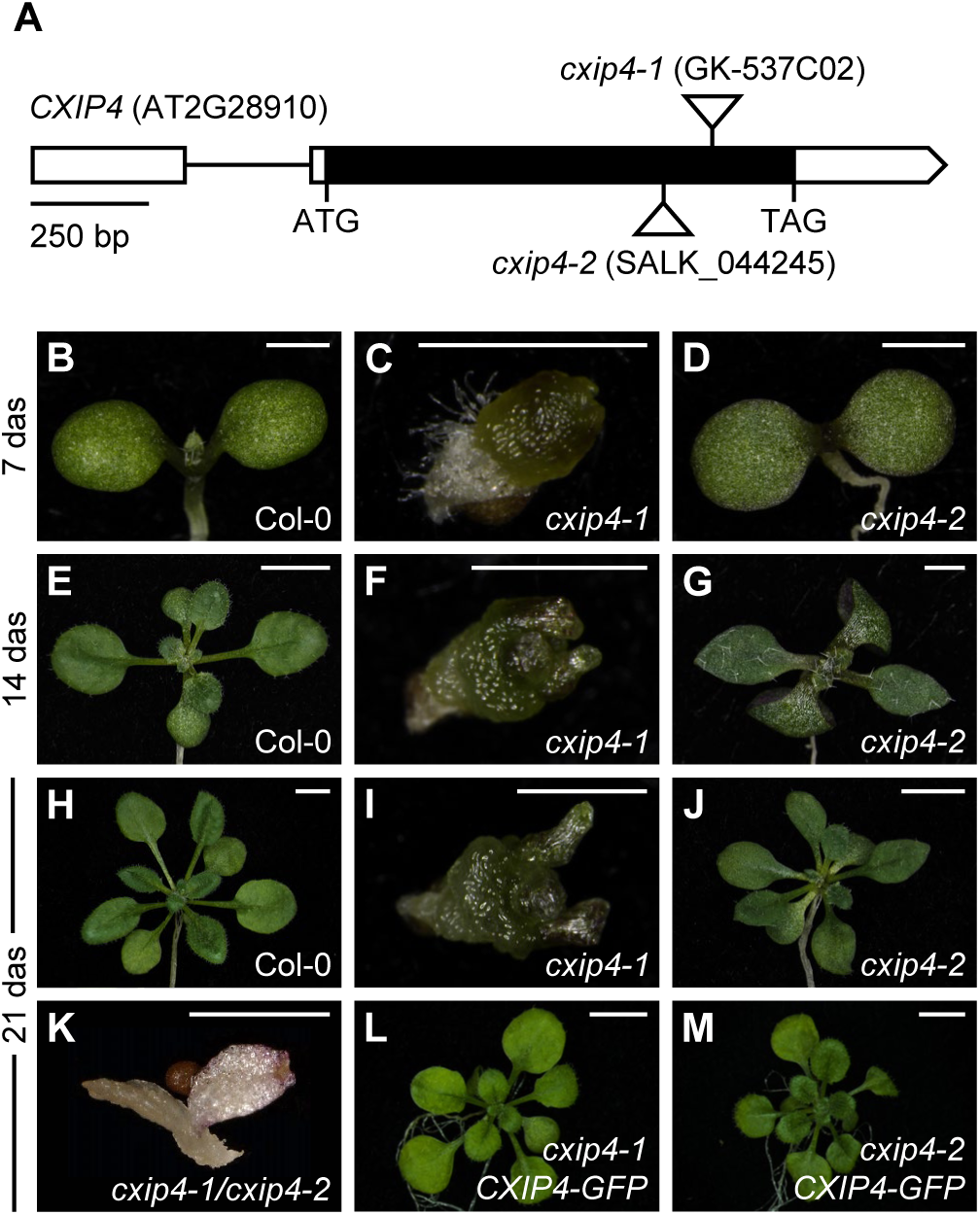
Structure of the *CXIP4* gene and developmental phenotypes of *cxip4* insertional alleles. (A) Schematic representation of the *CXIP4* gene, indicating the positions of its start (ATG) and stop (TAG) codons and the T-DNA insertions (triangles) in the *cxip4* alleles studied in this work. Boxes represent exons, with untranslated (UTRs) and coding regions shown in white and black, respectively. (B-M) Morphological vegetative phenotypes of (B, E, H) Col-0, (C, F, I) *cxip4-1*, (D, G, J) *cxip4-2*, (K) *cxip4-1/cxip4-2*, (L) *cxip4-1 CXIP4_pro_:CXIP4:GFP*, and (M) *cxip4-2 CXIP4_pro_:CXIP4:GFP* plants. Photographs were taken (B-D) 7, (E-G) 14, and (H-M) 21 days after stratification (das). Scale bars: (B, C, D, F, G, I, and K) 1 mm, and (E, H, J, L, and M) 5 mm.

To date, only a single functional analysis of the *CXIP4* gene in Arabidopsis has been performed, which was based on transgenic lines with β-estradiol inducible overexpression of full-length cDNA (Coego et al., 2014). To investigate the effects of the loss of function of *CXIP4*, we obtained two T-DNA insertional lines from public collections with disruptions in the coding region of AT2G28910: GK-537C02 and SALK_044245 (Fig. 1A). GK-537C02 seeds either did not germinate or produced phenotypically wild-type plants or callus-like seedlings with early arrested development (Fig. 1, B, C, E, F, H, and I). The phenotypically wild-type plants were found to be homozygous for the wild-type allele of *CXIP4* or hemizygous for the T-DNA insertion; siliques of the latter plants exhibited some light-green, immature seeds that contained early arrested embryos (Fig. 2, A to D). We sowed 3,626 additional seeds from hemizygous plants, finding that 15.6% exhibited a wrinkled appearance and failed to germinate (12.6%) or produced dark callus-like seedlings (2.9%), while the remaining 85.4% gave rise to phenotypically wild-type plants (Fig. 2, E and F). We genotyped 30 callus-like seedlings and 170 phenotypically wild-type plants. All aberrant seedlings were homozygous for the T-DNA insertion, whereas those with a wild-type phenotype were hemizygous or homozygous for the wild-type allele of *CXIP4*. We named the insertional allele of *CXIP4* carried by the GK-537C02 line *cxip4-1*. The *cxip4-1/cxip4-1* seedlings did not develop true leaves or any other recognizable organ (Fig.1, C, F, and I). The ungerminated seeds were assumed to also be homozygous for the *cxip4-1* allele, and we did not genotype them.

**Figure 2.**
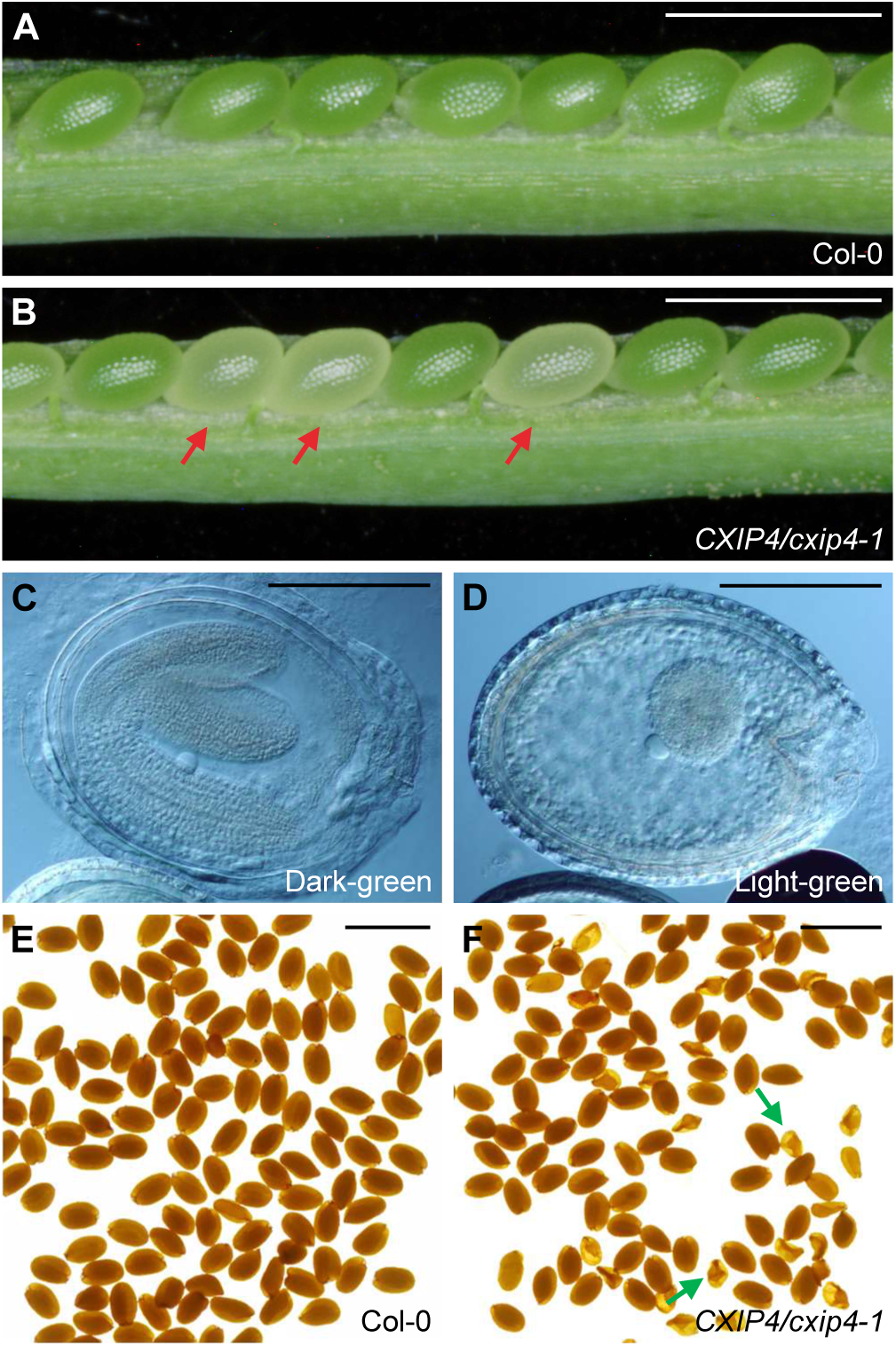
Developmental defects caused by the *cxip4-1* allele. (A, B) Dissected immature siliques from (A) Col-0 and (B) *CXIP4/cxip4-1* plants; the latter exhibit light-green seeds (red arrows). (C, D) Embryos in (C) dark-and (B) light-green seeds from siliques of *CXIP4/cxip4-1* plants. (E, F) Seeds from (E) Col-0 and (F) *CXIP4/cxip4-1* plants; the latter developed some wrinkled seeds (green arrows), whose genotype was found to be *cxip4-1/cxip4-1*. Photographs in (A-D) were taken 60 das. Scale bars: (A, B, E, and F) 1 mm, and (C, D) 100 µm.

Some seeds of the SALK_044245 line yielded plants with significantly slowed growth and defects in both vegetative and reproductive development (Fig. 1, D, G, and J; Fig. 3; Supplementary Fig. S2). Seven days after stratification (das), plants homozygous for the T-DNA insertion were smaller than Col-0 plants (8.8% of 979 seedlings; Fig. 1, B and D), and some had three cotyledons. At 14–21 das, the mutant plants showed pointed and reticulated leaves with a reduced density of trichomes, whose branching was not affected, and produced high levels of anthocyanins (Fig. 1, E, G, H, and J; Fig. 3, A to J). These plants flowered late, developed numerous shoots with reduced apical dominance, and remained green for more than 90 days (Fig. 3K and Supplementary Fig. S2). In addition, their flowers produced short siliques containing only a few seeds (Fig. 3, L and M). We named the insertional allele of *CXIP4* carried by the SALK_044245 line *cxip4-2*.

**Figure 3.**
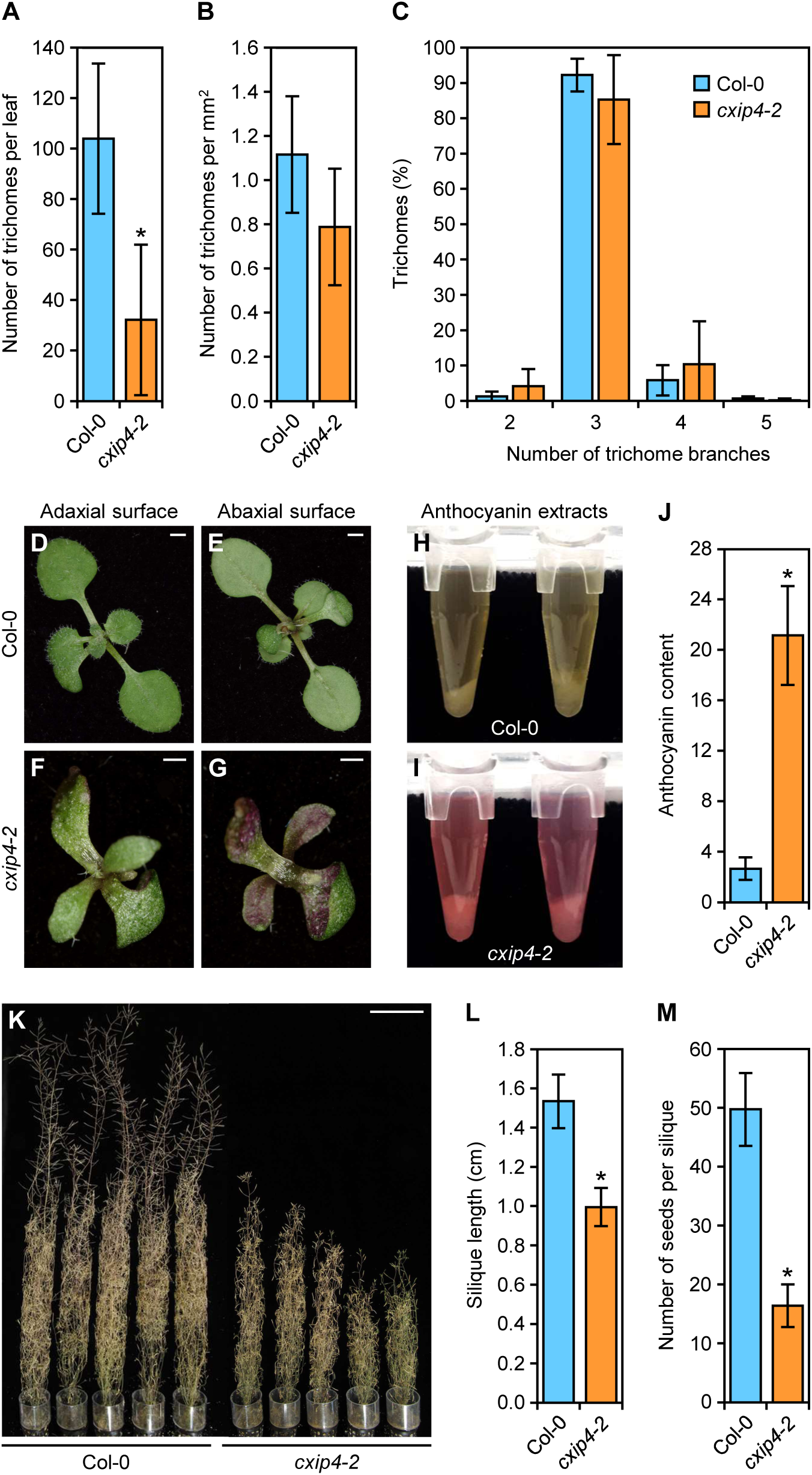
Pleiotropy of the morphological phenotype of *cxip4-2* plants. (A-C) Trichome density and branching in the third leaves of wild-type Col-0 and the *cxip4-2* mutant. Trichomes of ten leaves collected 21 (Col-0) or 29 (*cxip4-2*) das were analyzed per genotype. (D-J) Anthocyanin accumulation in the leaves of *cxip4-2*. (D-G) Adaxial and abaxial surfaces of rosettes of Col-0 and *cxip4-2* collected 12 das. (H, I) Anthocyanin extracts from Col-0 and *cxip4-2*, and (J) anthocyanin contents of ten biological replicates per genotype, each consisting of a rosette collected 12 das. (K) Adult Col-0 and *cxip4-2* plants collected 120 das. (L) Silique length, and (M) fertility of Col-0 and *cxip4-2* plants. Ten siliques collected from five plants per genotype were analyzed. Error bars in (A-C), (J), (L), and (M) represent standard deviations. Asterisks in (A), (J), (L), and (M) indicate values significantly different from the wild type in a Student’s *t*-test (\**p* < 0.0001). Scale bars: (D-G) 1 mm, and (K) 10 cm.

The T-DNA insertions in *cxip4-1* and *cxip4-2* are located close to each other (Fig. 1A), but their mutant phenotypes are quite different (Fig. 1, C, D, F, G, I, and J). Therefore, we analyzed the mRNAs produced by *cxip4-1* and *cxip4-2.* Both mRNAs were chimeric, as they included sequences from *CXIP4* and the T-DNA (Supplementary Fig. S3, A and B). The mRNA from *cxip4-1* was predicted to encode a mutant CXIP4-1 protein four aa longer than the wild-type CXIP4, with only 271 aa shared between the mutant and wild type. Translation of the chimeric *cxip4-2* mRNA would result in a protein of 240 aa missing part of the C-terminal region (Supplementary Figs. S1 and S3C). The aberrant CXIP4-1 protein appears to lack activity, but the truncated CXIP4-2 protein may be partially functional. These results might explain the stronger phenotype observed in *cxip4-1* plants compared to *cxip4-2*.

We carried out a complementation test by crossing *CXIP4/cxip4-1* to *cxip4-2/cxip4-2* plants. We obtained four *cxip4-1/cxip4-2* F_1_ plants from two different crosses, one of which displayed three cotyledons, as did some *cxip4-2/cxip4-2* plants. The four *cxip4-1/cxip4-2* heterozygous plants displayed a strong mutant phenotype more similar to that of *cxip4-1/cxip4-1* than *cxip4-2/cxip4-2* homozygotes, and their development was arrested after the emergence of cotyledons and one or two small leaves (Fig. 1K). These results confirm the allelism of *cxip4-1* and *cxip4-2*, and with the recessive nature of both, suggest that they are null and hypomorphic alleles of *CXIP4*, respectively.

To confirm that the disruption of *CXIP4* by the T-DNA insertions in *cxip4-1* and *cxip4-2* was the cause of their mutant phenotypes, we transferred a copy of the wild-type *CXIP4* gene into *CXIP4/cxip4-1* and *cxip4-2/cxip4-2* plants. Indeed, homozygous *cxip4-1 CXIP4_pro_:CXIP4:GFP* and *cxip4-2 CXIP4_pro_:CXIP4:GFP* plants were phenotypically wild type (Fig. 1, L and M).

### *CXIP4* is ubiquitously expressed in Arabidopsis

Cis-regulatory elements in Arabidopsis are usually located within the 500 base pair (bp) region upstream of the transcription start site (TSS), with the majority of these elements (86%) located proximal to the TSS (+1 bp position), spanning from −1,000 bp to +200 bp (Yu et al., 2016). AT2G28910 (*CXIP4*) and AT2G28920 are oriented with their respective 5′ ends toward each other, with their translation start codons 1,026 bp away from each other, sharing an intergenic region of 401 bp that could potentially contain a bidirectional promoter. No 5′-UTR is annotated for AT2G28920, but three different splice variants have been found for *CXIP4*, differing only in the length of the 5′-UTR, due to the AS of its single intron that interrupts this region. Therefore, we constructed two transgenes by cloning the 2,104- and 971-bp genomic regions upstream of the translation start codon of *CXIP4* into the pMDC164 vector, including 1,479 (*CXIP4_proI_*) or 346 (*CXIP4_proII_*) bp of the intergenic region, driving β−glucuronidase (GUS) expression.

We detected GUS activity in all organs of the *CXIP4_proI_:GUS* and *CXIP4_proII_:GUS* transgenic plants, with the highest activity in emerging leaves and the vasculature of cotyledons, leaves, and roots, especially in the root apex (Supplementary Fig. S4, A to D). We also detected GUS activity during embryogenesis in seeds at the green mature embryo stage, which is consistent with the early lethality found in *cxip4-1* (Supplementary Fig. S4E). We did not observe any differences between plants expressing the *CXIP4_proI_:GUS* (not shown) or *CXIP4_proII_:GUS* transgenes, suggesting that the 971 bp region upstream of the translation start codon of *CXIP4* contains all the regulatory elements needed for *CXIP4* expression. Indeed, the expression of the *CXIP4_proII_:CXIP4:GFP* transgene rescued the mutant phenotype of *cxip4-1* and *cxip4-2* plants (Fig. 1, L and M).

### Phylogenetic analysis of CXIP4 and other PTHR31437 family members

CXIP4 appears to be plant-specific, although it is classified as a member of the SREK1IP1 protein family (PTHR31437; Supplementary Table S1) according to the PANTHER database (https://www.pantherdb.org; Thomas et al., 2022). Human SREK1IP1 and Arabidopsis CXIP4 share only a 21-aa region, which harbors the 18-aa ZCCHC motif (Supplementary Fig. S5). It is of note that the ZCCHC motifs of CXIP4 and SREK1IP1 clearly grouped in a branch of the unrooted tree that we obtained from the alignment of 198 ZCCHCs from 110 proteins of yeast, human, and Arabidopsis (Aceituno-Valenzuela et al., 2020). This grouping reflects the remarkably high similarity (13 identical amino acids) of the ZCCHC motifs of these two proteins (Supplementary Fig. S5).

The PTHR31437 family, as described in the PANTHER database, comprises 96 members from the animal (SREK1IP1-like), plant (CXIP4-like), protozoa, and chromista kingdoms, with proteins of this family expected to be present in all organisms of these four kingdoms. While all animal and most plant genomes studied to date encode only one SREK1IP1/CXIP4 protein (such as Arabidopsis), some plants contain two to four co-orthologs (Supplementary Table S1).

We studied the amino acid sequences of the 96 proteins classified as members of the PTHR31437 family in the UniProtKB database. One of the two proteins of the ancestral angiosperm *Amborella trichopoda* and those from the chromistan *Phytophthora ramorum* and *Thalassiosira pseudonana* lacked the ZCCHC motif (Supplementary Table S1). We aligned the sequences of five ZCCHC-containing proteins from five distant organisms: *Chlamydomonas reinhardtii* (UniProtKB accession number A0A2K3DE08), *Arabidopsis thaliana* (Q84Y18), *Physcomitrium* (*Physcomitrella*) *patens* (A0A2K1K6W5), *Drosophila melanogaster* (Q9W3Z5), and *Homo sapiens* (Q8N9Q2). The alignment revealed high conservation only in the region containing the ZCCHC motif and in three isolated basic residues of K or R, located downstream of the ZCCHC motif and at the C-terminus of the proteins (Supplementary Fig. S5). We found five additional conserved amino acids, in addition to the three canonical C and one H of the ZCXCHC motif, which may be relevant for the functions of PTHR31437 family proteins. The alignment also showed that the *Drosophila melanogaster* and human proteins lack the N-terminal extension found in the CXIP4-like proteins of photosynthetic organisms. In addition, the ZCCHC motifs in *Drosophila melanogaster* and humans are located at the N-terminal regions of these proteins (Supplementary Fig. S5), as also detected in all metazoan SREK1IP1 orthologs, whose lengths were half those of plant CXIP4-like proteins (Supplementary Table S1).

We also aligned the sequences of putative CXIP4-like proteins from different angiosperm lineages, selecting species encoding a single ortholog: *Hordeum vulgare*, *Arabidopsis thaliana*, *Capsicum annuum*, *Vitis vinifera*, and *Populus trichocarpa*. This second alignment revealed higher conservation than that shown in Supplementary Fig. S5, as expected, including a 48-aa region containing the ZCCHC motif (Supplementary Fig. S6) and the full conservation of the N-terminal 39 aa, which are absent in animals and in the moss *Physcomitrium patens*, and partially conserved in the single-celled green alga *Chlamydomonas reinhardtii* (Supplementary Figs. S5 and S6). In fact, when we performed a BLASTP search using that 39-aa conserved sequence as a query, we only found the exact sequence in CXIP4 proteins from divergent land plant lineages of five major clades: bryophytes (such as *Physcomitrium patens*), lycophytes, monilophytes, gymnosperms, and angiosperms (such as Arabidopsis).

### CXIP4 is a nucleoplasmic protein

When transiently expressed in tobacco (*Nicotiana tabacum*) plants, Arabidopsis CXIP4 localized to the nucleus in a diffuse pattern; however, when expressed in yeast and tobacco BY-2 cells, it was also found in the cytoplasm, forming discrete spots that did not correspond to mitochondria (Cheng et al., 2004). More recently, Arabidopsis CXIP4 was found to exclusively localize to the nuclei of chickpea (*Cicer arietinum*) protoplasts (Cheng and Nakata, 2020).

As an initial assessment of the localization of Arabidopsis CXIP4, we used the MULocDeep web server (https://www.mu-loc.org/; Jiang et al., 2021; Jiang et al., 2023), which predicts the localization of any eukaryotic protein in 44 suborganellar compartments based on its primary sequence. We included human SREK1IP1 in the analysis. MULocDeep predicted that both proteins localize to the nucleus, with scores of 0.840 and 0.996, respectively (Supplementary Fig. S7A). Within the nucleus, both proteins are predicted to predominantly localize to the nucleoplasm, nucleolus, and nuclear speckles (Supplementary Fig. S7B), which are known to be enriched in pre-mRNA splicing factors, snRNAs, and polyadenylated [poly(A)^+^] RNAs (Belmont, 2022).

Using the LOCALIZER (https://localizer.csiro.au/; Sperschneider et al., 2017) and NoD (https://www.compbio.dundee.ac.uk/www-nod/index.jsp; Scott et al., 2010; Scott et al., 2011) web tools, we predicted nuclear and nucleolar localization signals (NLSs and NoLSs, respectively) in the C-terminal half of CXIP4. These NLSs and NoLSs signals were identified in two or three separate stretches, respectively, each composed of positively charged aa, mainly K and R (Supplementary Fig. S1).

To determine the localization of Arabidopsis CXIP4, we analyzed Col-0 plants carrying the *CXIP4_pro:_CXIP4:GFP* transgene, which successfully rescued the mutant phenotypes of *cxip4-1* and *cxip4-2* plants (Fig. 1, L and M), revealing that the CXIP4-GFP fusion protein is functionally equivalent to the endogenous CXIP4. The CXIP4-GFP fusion protein appeared in a speckled pattern within the nucleus (Fig. 4), which may correspond to nuclear speckles, as predicted by MULocDeep (Supplementary Fig. S7B).

**Figure 4.**
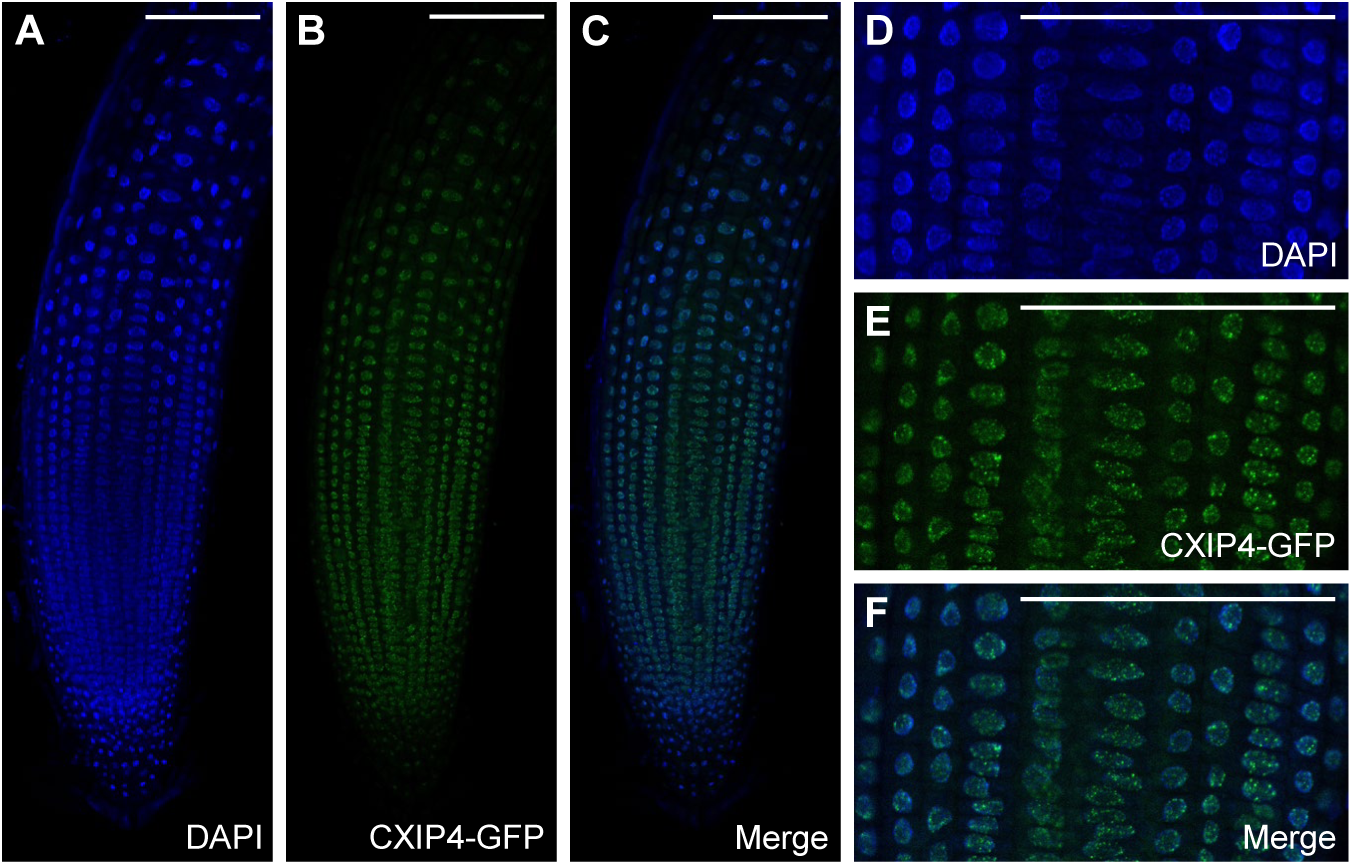
Subcellular localization of CXIP4 in Arabidopsis roots collected 5 das. (A-F) Confocal laser-scanning micrographs of Col-0 plants homozygous for the *CXIP4_pro_:CXIP4:GFP* transgene. Fluorescent signals correspond to (A, D) 4′,6-diamidino-2-phenylindole (DAPI) nuclear staining (in blue), (B, E) CXIP4-GFP (in green), and (C, F) their overlap. Scale bars: 100 μm.

### Human SREK1IP1 interacts with splicing factors that participate in AS

In GeneCards of the Human Gene Database (https://www.genecards.org/), SREK1IP1 is described as “Gene Ontology (GO) annotations related to this gene include nucleic acid binding”, and in UniProtKB (https://www.uniprot.org/uniprotkb) as “Possible splicing regulator involved in the control of cellular survival”. Due to the dearth of functional information on PTHR31437 family members, we carried out searches in several databases. We found relevant information on human SREK1IP1 in the HitPredict database (http://www.hitpredict.org), which compiles physical protein–protein interactions from six different databases and 124 species, all experimentally identified in both small-scale and high-throughput assays (López et al., 2015). In HitPredict, we found an interaction with NKAP, which was obtained through an affinity purification mass spectrometry (AP-MS) high-throughput assay (Huttlin et al., 2017), and with several splicing factors (Supplementary Table S2). These results confirm the conservation of the interactions of SREK1IP1/CXIP4 with NKAP/MAS2 between humans and Arabidopsis, and they support a role for human SREK1IP1 in pre-mRNA splicing.

### *cxip4-2* globally affects the expression of genes involved in pathogen defense

To gain insight into the biological processes affected in the *cxip4* mutants, we performed RNA-seq analysis using poly(A)^+^ RNAs from Col-0 and *cxip4-2* plants collected 14 das. We identified 2,160 deregulated genes in *cxip4-2* compared to the wild type, with a similar number of up-(1,094) and downregulated (1,066) genes (Fig. 5A; Supplementary Data Set S1). *CXIP4* was the tenth most downregulated gene, suggesting that the T-DNA insertion disrupting *CXIP4* in the *cxip4-2* mutant strongly reduces its transcription (Fig. 5, A and B; Supplementary Data Set S1). However, compared with the early lethality exhibited by *cxip4-1*, the low expression of *CXIP4* in *cxip4-2* suggests that it is sufficient for viability, but strongly compromises fertility, in addition to causing a highly pleiotropic developmental phenotype (Fig. 1, B to J; Fig. 2; Fig. 3; Supplementary Fig. S2).

**Figure 5.**
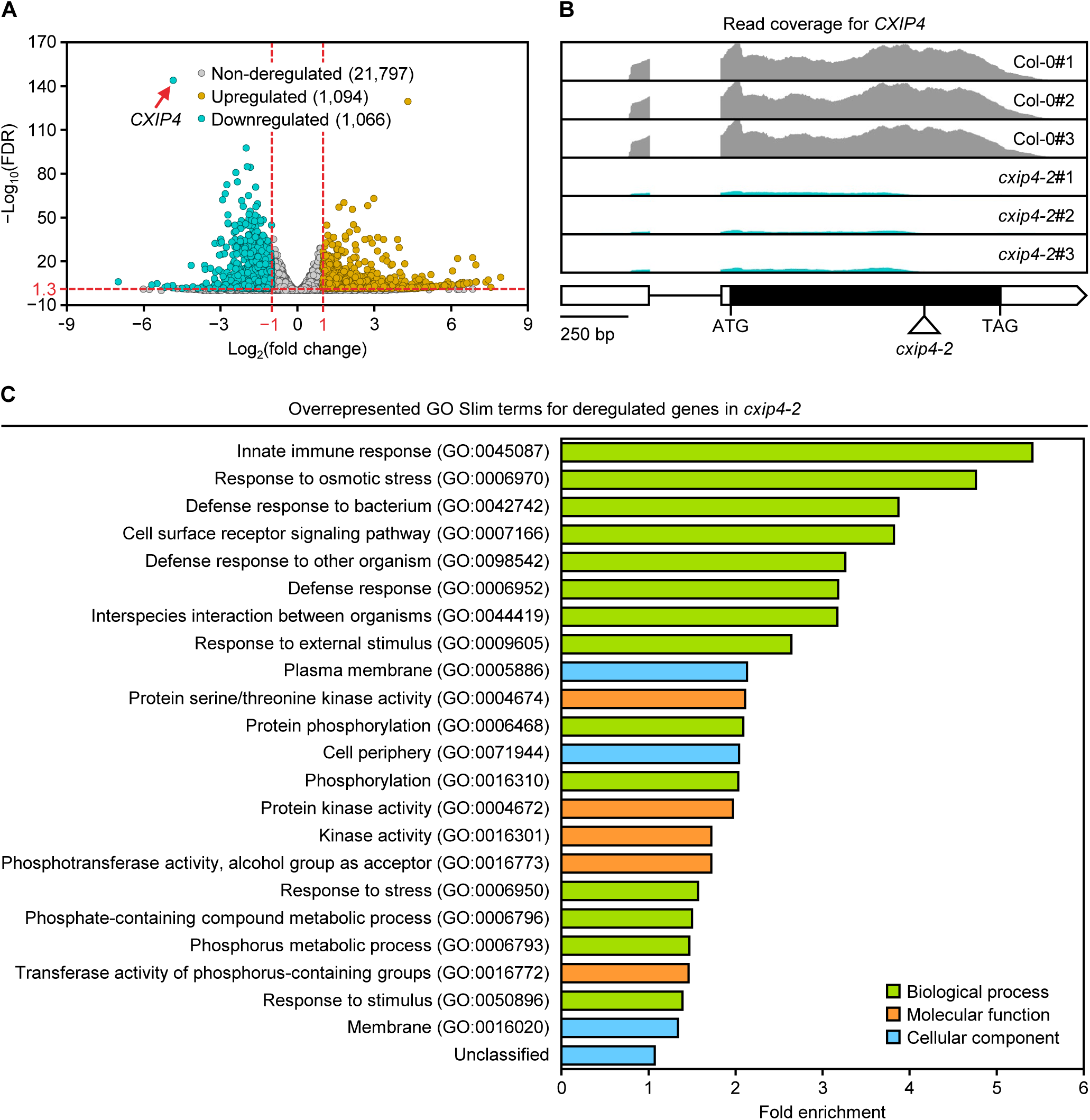
Differential gene expression analysis between Col-0 and *cxip4-2* plants. (A) Volcano plot of differentially expressed genes in *cxip4-2* compared to Col-0 plants, collected 14 das. Upregulated genes (fold change ≥2) and downregulated genes (fold change ≤2) and with FDR <0.05 (Benjamini-Hochberg false discovery rate) are shown as yellow and cyan dots, respectively; non-deregulated genes appear as gray dots. Horizontal and vertical red dashed lines mark the cutoffs of the negative decimal logarithm of FDR and binary logarithm of fold changes. The dot marked with a red arrow represents *CXIP4*, which was downregulated in *cxip4-2*, as expected. (B) Plot of aligned reads for *CXIP4* in Col-0 and *cxip4-2*, obtained with IGV software (http://software.broadinstitute.org/software/igv/). Gene structure is represented as described in the Figure 1 legend. (C) Bar plot representation of the overrepresented GO Slim terms for deregulated genes in *cxip4-2.* These terms were identified through statistical overrepresentation tests using PANTHER (https://pantherdb.org/) GO Slim annotation sets for biological processes, molecular functions, and cellular components (fold enrichment >1 and FDR <0.05). Redundant terms, with an identical set of genes to others that were already included, are not plotted. For a separate analysis of upregulated and downregulated genes in *cxip4-2*, see Supplementary Data Set S1.

Gene ontology (GO) enrichment analysis of the deregulated genes in *cxip4-2* plants revealed an overrepresentation of terms primarily associated with pathogen defense (Fig. 5C). In a separate analysis of downregulated genes, we also found an overrepresentation of terms related to indole glucosinolate, camalexin, and phytoalexin metabolism, as well as salicylic acid synthesis and perception, among others. By contrast, upregulated genes were mainly associated with responses to sulfur starvation and other stress conditions, as well as flavonoid, glucosinolate, and jasmonic acid metabolism (Supplementary Data Set S1).

Taken together, these results suggest that the deregulation of *CXIP4* causes a global defense response to biotic and abiotic stress. Interestingly, the upregulated genes were also enriched in DNA-binding transcription factor activity, including 80 out of 1,208 genes in the reference database, mainly with the C_2_H_2_ zinc finger (8 genes) and homeobox (32 genes) DNA-binding domains (Supplementary Data Set S1).

### Several upregulated genes in *cxip4-2* may explain its mutant phenotype

The *cxip4-2* mutant is viable but exhibits a highly pleiotropic phenotype, suggesting that many developmental pathways or key genes are downregulated in the mutant. Therefore, we inspected the RNA-seq data, looking for specific deregulated genes that may explain some of the phenotypes exhibited by *cxip4-2* plants. Interestingly, *GLABROUS1* (*GL1*), encoding a positive regulator of trichome development (Larkin et al., 1994), was downregulated, and *ENHANCER OF TRY AND CPC2* (*ETC2*), which encodes a MYB-like transcription factor family member that negatively regulates trichome development (Kirik et al., 2004), was upregulated (Supplementary Data Set S1). These results could explain the low density of trichomes found in *cxip4-2* leaves, whereas trichome branching, which is not controlled by these genes, remained unaffected (Fig. 3, A to C).

Among the upregulated genes in *cxip4-2*, we also identified *ENHANCED RNA INTERFERENCE-1-LIKE-1* (*ERIL1*; Supplementary Data Set S1), which encodes a ribonuclease H-like protein involved in the processing of chloroplast pre-rRNAs (Mermigka et al., 2016). Transgenic plants overexpressing *ERIL1* are defective in chloroplast rRNA maturation, resulting in the accumulation of several rRNA biogenesis intermediates. These precursors include a 1.7-kb transcript that may correspond to the 16S rRNA containing the intergenic region that separates this gene from the *trnl* gene, as well as the 23S-4.5S dicistronic precursor (3.2 kb), and the 2.4 and 2.9 kb incompletely processed 23S rRNAs (Mermigka et al., 2016; Supplementary Fig. S8A). The upregulation of *ERIL1* in *cxip4-2* could explain the presence of two additional peaks in the profiles of total RNA samples, which we obtained before their use in the RNA-seq assay. Both peaks are located between those corresponding to the cytosolic 18S and 25S rRNAs, whose sizes in Arabidopsis are approximately 1.8 and 3.4 kb, respectively, and could correspond to the incompletely processed chloroplast 23S rRNAs mentioned above (Supplementary Fig. S8B).

*PRODUCTION OF ANTHOCYANIN PIGMENT 1* (*PAP1*) was also found among the upregulated genes in *cxip4-2* (Supplementary Data Set S1). PAP1 is a transcription factor that directly induces the transcription of genes involved in anthocyanin biosynthesis. *PAP1*-overexpressing plants accumulate high amounts of anthocyanins when grown in the presence of high sucrose concentrations, which causes stress (Teng et al., 2005). The upregulation of *PAP1*, along with the enriched GO terms “anthocyanin-containing compound biosynthetic process” and “flavonoid metabolic process” (Supplementary Data Set S1), may also account for the high anthocyanin levels observed in the *cxip4-2* mutant (Fig. 3, D to J), suggesting that *cxip4-2* plants are constitutively stressed.

We did not find enriched GO terms related to the UPR, as might have been anticipated given the resistance to DTT and TM observed in plants overexpressing *CXIP4* (Hossain et al., 2016). Nevertheless, among the downregulated genes in *cxip4-2* plants was AT1G42990, which encodes the BASIC REGION/LEUCINE ZIPPER MOTIF 60 (bZIP60) transcription factor. bZIP60 activates the transcription of key genes involved in UPR signaling (Iwata and Koizumi, 2005).

### Pre-mRNA splicing is defective in *cxip4-2* plants

As previously mentioned, the GO annotations of human SREK1IP1 and its interactions associate this protein with pre-mRNA splicing. Thus, we analyzed the effects of *cxip4-2* on pre-mRNA splicing, finding 939 differential AS events in *cxip4-2* plants compared to Col-0. The most frequent were intron retentions (IR), with 543 events (57.8% of the total differential AS events), mainly resulting in the retention of the alternatively spliced intron (Fig. 6A; Supplementary Data Set S2).

**Figure 6.**
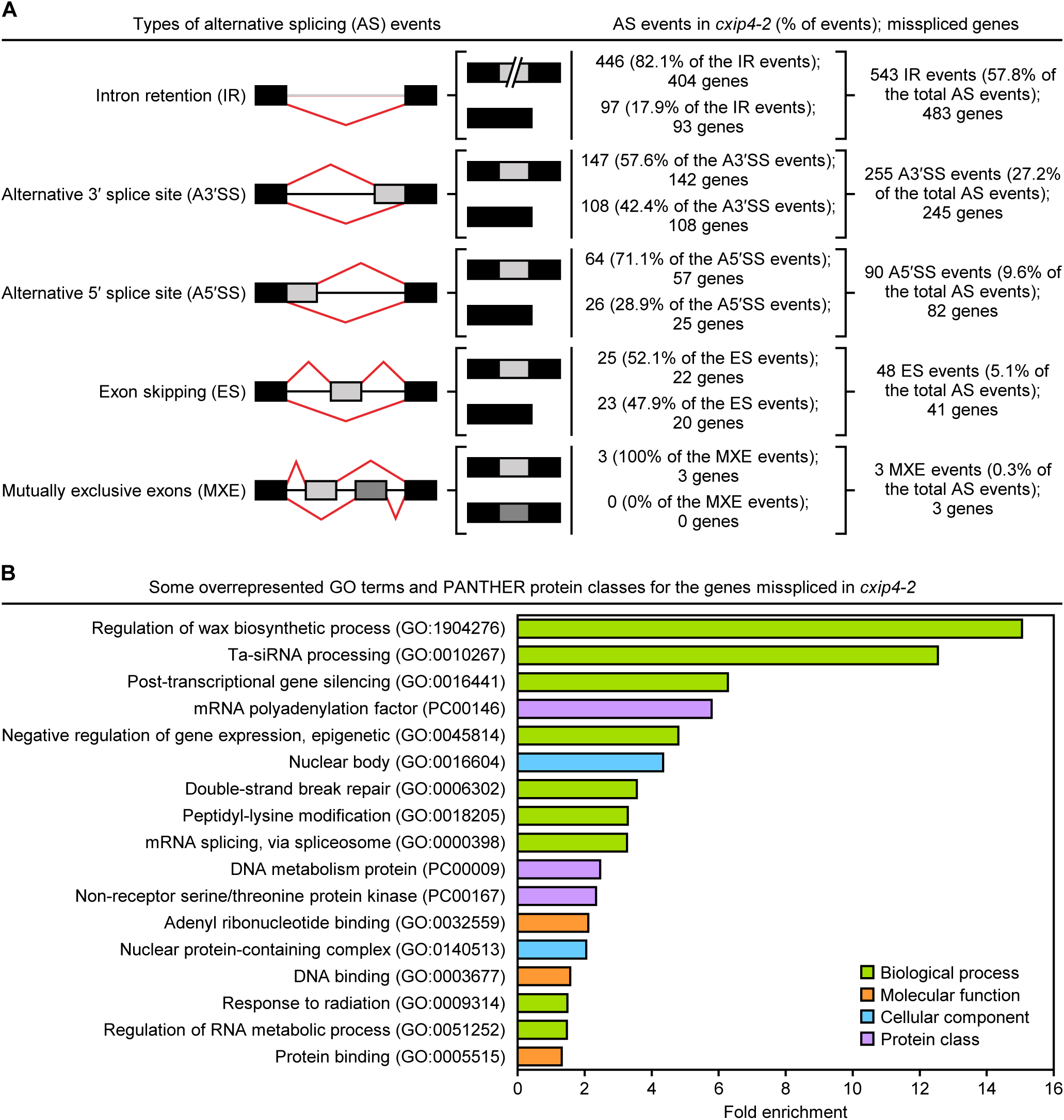
Analysis of differential alternative splicing between Col-0 and *cxip4-2* plants. (A) Summary of differential alternative splicing (AS) events and misspliced genes identified in *cxip4-2* compared to Col-0 at 18 das. Schematic representations depict five different types of AS events, with black or gray rectangles and lines representing constitutive or alternatively spliced exons and introns, respectively, and red lines marking the splice junctions. (B) Bar plot of overrepresented GO terms and PANTHER protein classes for the misspliced genes in *cxip4-2.* These terms were identified as described in the Figure 5 legend, but using GO Complete and PANTHER protein class annotation sets. Only the most specific hierarchically organized GO terms and PANTHER protein classes were plotted; for a complete list, see Supplementary Data Set S2.

The second most frequent differential AS events identified in *cxip4-2* plants compared to Col-0 were alternative 3′ splice site (A3′SS) events, totaling 255 (27.2%). Using Integrative Genomics Viewer (IGV; Robinson et al., 2011), we determined that 85 out of these 255 events (33.3% of the differential A3′SS events) involved tandem 3′SSs (acceptor sites) with NAGNAG sequences (Supplementary Data Set S2), in which A3′SSs were exactly three nucleotides apart from each other (in-frame); their alternative usage would result in proteins with one single amino acid of difference, which might not impact their function. The tandem 3′SS motif NAGNAG is overrepresented in genes encoding RNA-recognition motifs (RRMs), including SR proteins (Schindler et al., 2008). However, we did not find enrichment in any protein class for these 85 A3′SS events, which included genes related to pre-mRNA splicing.

### Subsets of misspliced genes in *cxip4-2* are related to different steps of mRNA metabolism

We analyzed the overrepresented GO terms for misspliced genes in *cxip4-2*, finding that they were mainly related to the regulation of cuticular wax biosynthesis, RNA-mediated gene silencing, mRNA splicing, chromatin remodeling, and DNA repair, among others (Fig. 6B; Supplementary Data Set S2). Five out of 15 genes in the reference database were categorized under the “ta-siRNA processing” term: *SUPPRESSOR OF GENE SILENCING 1* (*SGS1*), *SGS2* (also known as *RNA DEPENDENT RNA POLYMERASE 6* [*RDR6*], and *SGS3*, with differential IR events; and *DICER-LIKE 3* (*DCL3*) and *DOUBLE-STRANDED-RNA-BINDING PROTEIN 4* (*DRB4*), with differential A3′SS events. These five genes, whose expression was not deregulated in *cxip4-2* plants, were also classified under the “siRNA processing” term (Supplementary Data Set S2). Other overrepresented GO terms with several overlapping genes included “post-transcriptional gene silencing”, “regulatory ncRNA processing”, “RNA-mediated gene silencing”, and “negative regulation of gene expression, epigenetic” (Supplementary Data Set S2).

We also analyzed the PANTHER protein classes that were overrepresented among the misspliced genes in *cxip4-2*, which included the “mRNA polyadenylation factor” class, with six out of 39 genes in the reference database. These six genes encode NOT1 (the scaffold protein of the CCR4-NOT complex), the deadenylases CATABOLITE REPRESSOR 4C (CCR4C) and CCR4D, POLY(A) POLYMERASE 2 (PAPS2) and PAPS4, and MATERNAL EFFECT EMBRYO ARREST 44 (MEE44). The differential AS events included IR events in *CCR4D*, *PAPS2*, *PAPS4*, and *MEE44*, and A3′SS events in *NOT1* and *CCR4C*, although these genes were not deregulated in *cxip4-2*. These proteins are thought to function in polyadenylation-assisted RNA degradation through the exonucleolytic cleavage of the poly(A) tail, thereby contributing to mRNA export, nuclear quality control, and translation (Passmore and Coller, 2022).

## DISCUSSION

### *CXIP4* is an essential gene whose partial loss of function triggers pronounced pleiotropic changes

No mutational analyses of *CXIP4-like* genes have been described, and only a single functional analysis of the *CXIP4* gene in Arabidopsis has been performed, which used transgenic lines with β-estradiol-inducible overexpression of full-length cDNAs (Coego et al., 2014). Subsequent analysis of these lines revealed the resistance of *CXIP4* overexpressing plants to DTT and TM, suggesting a role for CXIP4 in the UPR (Hossain et al., 2016). However, the overexpression of a gene can cause artifacts, which does not occur in loss-of-function analysis. Moreover, loss-of-function analysis is usually more informative in the absence of functional redundancy. This would be the case for *CXIP4*, since it is a single-copy gene in Arabidopsis (Supplementary Table S1).

The functional analysis of two T-DNA insertional lines of *CXIP4* harboring alleles with different degrees of loss of function allowed us to conclude that *CXIP4* is an essential gene for embryogenesis but is also required for postembryonic development (Figs. 1 to 3; Supplementary Fig. S2). *cxip4-1* is likely a null allele of *CXIP4* that mainly causes early embryonic lethality, and the few embryos that escape this lethality produce callus-like plants with arrested development (Fig. 1, A, B, C, E, F, H, I, and L; Fig. 2). The c*xip4-2* allele, which is characterized by significantly reduced *CXIP4* expression, as identified by RNA-seq analysis (Fig. 4, A and B), is fully viable yet has a striking pleiotropic phenotype when homozygous (Fig. 1, A, B, D, E, G, H, J, and M; Fig. 3; Supplementary Fig. S2). This phenotype includes severe growth retardation leading to delayed flowering, abnormal cotyledon and leaf morphology characterized by sparse trichome density, the loss of apical dominance, and markedly reduced fertility (Fig. 3). Our analysis revealed the deregulation of several key genes, predominantly encoding transcription factors, in *cxip4-2* plants, providing insights into the underlying mechanisms of its pleiotropic phenotype (Supplementary Data Set S1). Furthermore, our findings suggest that the C-terminal region of CXIP4, which is absent from the CXIP4-2 mutant protein, is dispensable for plant survival but crucial for proper development (Supplementary Fig. S1). The pleiotropy exhibited by *cxip4-2* and the embryonic lethality of *cxip4-1* match the broad GUS activity detected in all tissues, including embryos, where expression was very high (Supplementary Fig. S4).

Besides their aberrant morphological phenotype, *cxip4-2* plants accumulate high levels of anthocyanins, suggesting that they are constitutively stressed. This is consistent with the finding that the deregulated genes identified by RNA-seq analysis were enriched in GO terms involved in global defense responses (Fig. 3, D to J; Fig. 5C). These results also agree with the recently proposed function of TaCAXIP4, one of the four putative co-orthologs of CXIP4 in wheat (Supplementary Table S1), in the calcium-mediated plant immune response against pathogens. The role of TaCAXIP4 in this defense response results from its interaction with TaHRC, a histidine-rich calcium-binding nuclear protein conferring susceptibility to Fusarium head blight or scab, a destructive disease in wheat worldwide caused by *Fusarium graminearum* (Su et al., 2019). It has been proposed that the interaction between TaHRC and TaCAXIP4 leads to the hijacking of TaCAXIP4, resulting in the suppression of calcium-mediated immune responses (Chen et al., 2022).

### Arabidopsis CXIP4 and human SREK1IP1 are highly divergent orthologs

The PANTHER database (https://www.pantherdb.org/) contains the complete proteomes of 143 organisms, including model organisms and others used in biomedical and biotechnological research. A search in PANTHER revealed that metazoan SREK1IP1-like and plant CXIP4-like proteins form the PTHR31437 family, but their sequences are highly divergent. In fact, they are classified as LDOs (least diverged orthologs), which are the most nearly ‘equivalent’ gene pairs between different organisms based on phylogenetic analysis. Metazoan SREK1IP1-like proteins are approximately half the length of plant CXIP4-like proteins, sharing only some amino acids from the ZCCHC motif (Supplementary Table S1 and Supplementary Fig. S5). However, it is of note that the ZCCHC motifs of Arabidopsis CXIP4 and human SREK1IP1, which share 13 to the 18 aa, clearly grouped in the same branch of an unrooted tree generated by phylogenetic analysis based on sequence alignment of 198 ZCCHC motifs from 110 ZCCHC-containing proteins in yeast (7), Arabidopsis (69), and humans (34) that we identified in a systematic search (Aceituno-Valenzuela et al., 2020). This grouping was likely due to the high number of conserved amino acids, in addition to the three canonical cysteines (C) and one histidine (H) that define the ZCCHC motif.

Despite the low similarity between Arabidopsis CXIP4 and human SREK1IP1 outside the ZCCHC domain, their subcellular and suborganellar localizations appear to be highly similar, as predicted by MULocDeep (Supplementary Fig. S7). MULocDeep uses a machine learning method based on primary sequence information (https://www.mu-loc.org/; Jiang et al., 2021; Jiang et al., 2023) to identify common signals or patterns in two proteins. In fact, the NLS and NoLS signals were present in the C-terminal half of CXIP4, which is rich in basic aa, similar to human SREK1IP1. These findings support the notion that these proteins share similar functions.

### The specific categories of misspliced genes in *cxip4-2*, rather than their number, may explain its mutant phenotype

The *prp8-7* hypomorphic allele of *PRE-MRNA PROCESSING 8* (*PRP8*), encoding a central factor of the spliceosome, causes global missplicing, as revealed by its 8,124 increased IR events compared to Col-0 (Sasaki et al., 2015), which is 18 times higher than in the *cxip4-2* plants (446 increased IR events; Fig. 6A; Supplementary Data Set S2). However, the morphological phenotype of *cxip4-2* is much stronger than that of *prp8-7* with both mutants exhibiting a partial loss-of-function of the corresponding essential genes (*CXIP4* and *PRP8*). Therefore, the phenotype of *cxip4-2* plants is not caused by the number of missplicing events, and it appears that CXIP4 does not play central roles in general splicing but suggests that it functions in other processes.

We also found that most of the detected differential AS events do not deregulate the expression of the affected genes, as is the case with the *prp8a-14* mutant of the *PRP8* gene (Llinas et al., 2022). It is possible that the missplicing events that we found in *cxip4-2* plants cause a reduction of functional mRNAs and alter the relative concentrations of the resulting protein isoforms, which may have diverse functional consequences. However, the overrepresentation of specific GO terms, mainly “gene silencing by DNA methylation” or the protein class “mRNA polyadenylation factor” among the misspliced genes in *cxip4-2* plants (Supplementary Data Set S2) suggests a role for CXIP4 in transcriptional and post-transcriptional regulation of gene expression, which should be studied further.

## MATERIALS AND METHODS

### Plant material, growth conditions and genotyping

All *Arabidopsis thaliana* (L) Heynh. lines used in this work were in the Columbia-0 (Col-0) genetic background. The Col-0 wild type and the *cxip4-1* (GABI_537C02) and *cxip4-2* (SALK_044245) T-DNA insertional mutants (Alonso et al., 2003; Kleinboelting et al., 2012) were obtained from the Nottingham Arabidopsis Stock Center (NASC; Nottingham, United Kingdom) and propagated in our laboratory for further analysis.

Seed sterilization and sowing, plant culture, and crosses were performed as previously described (Ponce et al., 1998; Berná et al., 1999), except that plant agar was replaced with 6 g·L^−1^ of Gelrite (Duchefa Biochemie). When required, culture media were supplemented with hygromycin (15 µg·mL^−1^) and kanamycin (50 µg·mL^−1^).

To genotype the *cxip4* alleles, genomic DNA was extracted from the samples as described in Ponce et al. (2006), and the presence of the T-DNA insertions in *CXIP4* was verified by PCR amplification using the primers shown in Supplementary Table S3.

Most Sanger sequencing reactions and electrophoreses were carried out in our laboratory with ABI PRISM BigDye Terminator Cycle Sequencing kits and an ABI PRISM 3130xl Genetic Analyzer (Applied Biosystems). Some sequencing reactions were carried out at Stab Vida (Caparica, Portugal).

### RNA isolation, cDNA synthesis, and mutant mRNA analysis

Total RNA was isolated with TRIzol Reagent (Invitrogen) from three biological replicates per genotype, each consisting of 100 mg of rosettes collected 14 das. Prior to cDNA synthesis, the RNA was treated with TURBO DNase (Invitrogen). RT-PCR amplifications were performed as described in Casanova-Sáez et al. (2014). The mRNAs produced by the *cxip4-1* and *cxip4-2* mutant alleles were analyzed by Sanger sequencing of cDNAs, which were amplified using the primers shown in Supplementary Table S3.

### Construction of transgenes and analysis of transgenic lines

Transgenes were generated by Gateway cloning as described in Sánchez-García et al. (2015), using the pGEM-T Easy221 entry vector (provided by B. Scheres) and the pMDC107 and pMDC164 (Curtis and Grossniklaus, 2003) destination vectors.

To assess the temporal and spatial expression patterns of *CXIP4*, two constructs, *CXIP4_proI_:GUS* and *CXIP4_proII_:GUS*, were generated by PCR amplification of the 2,104-and 971-bp genomic regions upstream of the translation start codon of *CXIP4*. The PCR products were subcloned into the pMDC164 destination vector. GUS enzymatic activity was observed throughout the development of Col-0 plants homozygous for the *CXIP4_proI_:GUS* and *CXIP4_proII_:GUS* transgenes. All parts of the plants were imaged using bright-field microscopy under a Nikon D-Eclipse C1 laser-scanning confocal microscope. GUS staining was performed as previously described (Donnelly et al., 1999). The samples were incubated in X-Gluc buffer overnight at 37°C. The seed coat and embryo were separated before incubating the seeds in GUS staining solution.

To determine the subcellular localization of CXIP4, the *CXIP4 _proII_:CXIP4:GFP* construct was generated by PCR amplification of the 971-bp genomic region upstream of the translation start codon and the full-length coding sequence of *CXIP4* (without its stop codon), which was then subcloned into the pMDC107 destination vector.

The structural integrity of all constructs was verified by sequencing prior to their transfer into Arabidopsis plants by *Agrobacterium tumefaciens*-mediated transformation via the floral dip method (Clough and Bent, 1998). Primers used to obtain these constructs and for Sanger sequencing are described in Supplementary Table S3.

### Analysis of plant morphology and trichome density and branching

Rosette photographs were taken under a Nikon SMZ1500 stereomicroscope equipped with a Nikon DXM1200F or DS-Ri2 digital camera. For large rosettes, high-resolution images were obtained by taking multiple pictures of the same plant and assembling them with the Photomerge tool of Adobe Photoshop CS3 software. Adult plants were photographed with a Canon PowerShot SX200 IS camera.

Trichome density and branch number were determined using the third leaves of ten Col-0 and *cxip4-2* plants collected 21 and 29 das, respectively. The leaves were cleared with ethanol and chloral hydrate, mounted on slides, and photographed under a Leica DMRB microscope equipped with a Nikon DXM1200 digital camera (Nadi et al., 2023). The number of trichomes with two to five branches was scored for each leaf and expressed as a percentage of the total number of trichomes.

### Anthocyanin extraction and measurement

To quantify anthocyanin contents, ten biological replicates per genotype were used, each consisting of a rosette collected 12 das and weighed. Anthocyanins were extracted and homogenized in 45% (v/v) methanol and 5% (v/v) acetic acid buffer using glass beads and a MixerMill 400 (Retsch) automatic mixer. The samples were centrifuged twice for 5 min at 13,500 g to remove cellular debris. The absorbance of each clarified extract was measured spectrophotometrically at 530 and 657 nm for anthocyanin and contaminating chlorophyll in acidic extraction buffer, respectively (Mancinelli, 1990). Anthocyanin content was calculated as A_530_ − (0.25 · A_657_) · g^−1^ of rosette fresh weight, where 25% of the A_657_ reading was subtracted to account for chlorophyll degradation products.

### RNA-seq and analysis of alternative splicing

Total RNA was extracted using TRIzol Reagent (Invitrogen) from three biological replicates per genotype, each consisting of 40–80 mg of rosettes collected at 14 das. At least 6 µg of RNA was sent to Novogene (Cambridge, United Kingdom) for sequencing and subsequent gene expression profiling. An Agilent 2100 bioanalyzer with an RNA 6000 Nano Kit (Agilent Technologies) was used to assess RNA concentration and integrity. Total RNA samples underwent mRNA enrichment prior to random fragmentation. Libraries were produced with a NEBNext Ultra RNA Library Prep Kit for Illumina (New England Biolabs) and sequenced on an Illumina NovaSeq 6000 platform using a 2 × 150 bp run protocol. More than 50 million non-stranded 150 bp paired-end reads were generated from each library. All FASTQ files were submitted to the Sequence Read Archive (SRA) database of the National Center for Biotechnology Information (NCBI) under the BioProject accession number PRJNA1090680 (https://dataview.ncbi.nlm.nih.gov/object/PRJNA1090680?reviewer=7k932roo1316i2pp 926ljh9q2s).

Differential gene expression analysis was conducted by Novogene. Briefly, raw data were processed through in-house Perl scripts to remove low-quality reads and reads containing adapters or more than 10% uncertain nucleotides. Clean reads were aligned to the Arabidopsis Col-0 genome (TAIR10) with HISAT2 2.0.5 (Kim et al., 2019) and assembled with StringTie 1.3.3b (Pertea et al., 2015), using default parameters. The number of reads (counts) mapped to each gene was determined using FeatureCounts 1.5.0-p3 (Liao et al., 2014) with default parameters, giving the number of fragments per kilobase of transcript per million mapped reads (FPKM). The identification of differentially expressed genes between Col-0 and *cxip4-2* plants was performed using the DESeq2 R package 1.20.0, with the selection criteria set at a fold change > 2 and a Benjamini-Hochberg false discovery rate (FDR) < 0.05.

Differential AS analysis was carried out at the Bioinformatics for Genomics and Proteomics Unit of the Centro Nacional de Biotecnología (CNB, Madrid). Briefly, the quality and purity of the raw reads were determined using FastQC 0.11.9 (https://www.bioinformatics.babraham.ac.uk/projects/fastqc/) and FastQ Screen 0.14.1 (Wingett and Andrews, 2018), respectively. Reads were aligned to the Arabidopsis Col-0 genome (TAIR10) using STAR 2.7.10a (Dobin et al., 2013) with default parameters. Differential AS events between Col-0 and *cxip4-2* plants were assessed with rMATS-turbo 4.1.2 (Shen et al., 2014). Only those events with an FDR < 0.05, an absolute Delta (percent spliced-in, PSI) > 0.1 (10%), and an average number of reads covering the event > 5 were considered to be statistically significant.

Statistical tests for the overrepresented GO terms and PANTHER protein classes among the differentially expressed or spliced genes between Col-0 and *cxip4-2* plants were performed using Fisher’s Exact tests with FDR correction, using the GO database released 2023-01-05 (DOI: 10.5281/zenodo.7942786) and PANTHER 18.0 (released 2023-08-01; https://pantherdb.org/; Thomas et al., 2022). The selection criteria were set at a fold enrichment > 1 and an FDR < 0.05.

### Bioinformatic analysis using protein sequences

The multiple sequence alignments shown in Supplementary Figs. S5 and S6 were obtained using MUSCLE V3.8 (Edgar, 2004) from the EMBL-EBI bioinformatic web server (Madeira et al., 2022).

To assess the subcellular and suborganellar localizations of the Arabidopsis CXIP4 protein, the MULocDeep web server (https://www.mu-loc.org/; Jiang et al., 2021; Jiang et al., 2023) was utilized.

To predict the effect of eliminating part of the C-terminal half of CXIP4-2, the LOCALIZER (https://localizer.csiro.au/; Sperschneider et al., 2017) and NoD (https://www.compbio.dundee.ac.uk/www-nod/index.jsp; Scott et al., 2010; Scott et al., 2011) web tools were utilized.

### Accession numbers

Sequence data from this article can be found at The Arabidopsis Information Resource (TAIR; https://www.arabidopsis.org) under the accession number: *CXIP4* (AT2G28910).

## Supporting information

Supplementary Data Set S1

Supplementary Data Set S2

## ACKNOWLEDGEMENTS

The authors would like to thank J.A. García-Martín for the differential AS analyses; J. Castelló, D. Navarro, and M. Gomariz for their excellent technical assistance; and J.L. Micol for useful discussions and comments on the manuscript, as well as for the use of his facilities.

## AUTHOR CONTRIBUTIONS

M.R.P. obtained funding and conceived, designed, and supervised research. R.M.-P and R.S.-M. partially co-supervised this work. U.I.A.-V., S.F.-C., R.M.-P., R.S.-M., and A.R.-B. conducted the experiments. U.I.A.-V., S.F.-C., R.M.-P., and M.R.P. analyzed the data, created the figures, tables, and datasets and wrote the manuscript. All authors revised and approved the manuscript.

## FUNDING

This work was supported by grants from the Ministerio de Ciencia, Innovación y Universidades of Spain (PID2020-117125RB-I00 [MCI/AEI/FEDER, UE]) and the Generalitat Valenciana (PROMETEO CIPROM/2022/2) to M.R.P. R.M.-P. held a María Zambrano distinguished researcher contract funded by the Next Generation EU programs and administered by the Universidad Miguel Hernández. U.I.A.-V. held a predoctoral fellowship from the Generalitat Valenciana (GRISOLIAP/2016/134).

## SUPPLEMENTARY DATA

**Supplementary Figure S1.** Localization of the ZCCHC motif, R-rich region, and nuclear and nucleolar localization signals in the predicted wild-type CXIP4 and mutant CXIP4-1 and CXIP4-2 proteins. The ZCCHC motif is highlighted in green, with the conserved cysteine and histidine residues (CX_2_CX_4_HX_4_C) shown in red letters. The R-rich region, as annotated in UniProtKB (https://www.uniprot.org/uniprotkb), is underlined. Nuclear localization signals (NLSs) are denoted with red letters, and nucleolar localization signals (NoLSs) are highlighted in yellow. The 65 aa at the C-terminus of CXIP4-1 (blue letters) replace the 61 aa of the corresponding region of CXIP4. CXIP4-2 contains 240 aa, with only the last three absent from CXIP4 (blue letters).

**Supplementary Figure S2.** Reproductive development of Col-0 and *cxip4-2* plants over time. Photographs were taken (A) 54, (B) 64, (C) 74, and (D) 84 das. Scale bars: 10 cm.

**Supplementary Figure S3.** Molecular effects of the T-DNA insertions in the *CXIP4* transcript. (A) Schematic representation of the *CXIP4* gene, as described in the Figure 1 legend, showing the positions of the primers used to analyze the effects of the T-DNA insertions (Supplementary Table S3). A segment of the gene is highlighted in blue and enlarged to better visualize such effects. Primers were not drawn to scale and are labeled as F (AT2G28910-F), LB1 (o8409), and LB2 (LBb1.3). (B) Agarose gel showing the PCR products obtained using cDNA and genomic DNA (gDNA) from *cxip4-1* and *cxip4-2* plants as templates and the F + LB1 and F + LB2 primer pairs. (C) Sequences of the chimeric *cxip4-1* and *cxip4-2* cDNAs and their predicted translation products CXIP4-1 and CXIP4-2. Nucleotide sequences corresponding to T-DNAs and the aa they are predicted to encode are highlighted in red. Asterisks indicate translation stop codons.

**Supplementary Figure S4.** Spatial expression of *CXIP4* during Arabidopsis development. GUS activity in Col-0 *CXIP4_proII_:GUS* transgenic plants carrying the short region of the *CXIP4* promoter during (A, B) vegetative and (C-E) reproductive development. GUS staining was performed using (A, B) whole plants collected (A) 7 and (B) 14 das; (C, D) inflorescences, siliques, and anthers from 50 das-old plants; and (E) embryos dissected from green mature seeds of 52 das-old plants. Scale bars: (A-D) 2 mm, and (E) 100 µm. Equivalent GUS staining results were obtained from Col-0 *CXIP4_proI_:GUS* plants.

**Supplementary Figure S5.** Sequence conservation among putative eukaryotic CXIP4 orthologs. Alignment of full-length putative CXIP4 orthologs from *Chlamydomonas reinhardtii* (UniProtKB accession number A0A2K3DE08), *Arabidopsis thaliana* (Q84Y18), *Physcomitrium patens* (A0A2K1K6W5), *Drosophila melanogaster* (Q9W3Z5), and *Homo sapiens* (Q8N9Q2). Asterisks and dots in the consensus line indicate identical (also shaded in black) and conserved residues, respectively. The consensus sequence of the ZCCHC motif is underlined in red.

**Supplementary Figure S6.** Sequence conservation among CXIP4 orthologs in angiosperms. Alignment of full-length CXIP4 orthologs from different angiosperm lineages: *Hordeum vulgare* (UniProtKB accession number A0A287R5P6) representing Poales, *Arabidopsis thaliana* (Q84Y18) for Malvids, *Capsicum annuum* (A0A2G3ANM6) for Asterids, *Vitis vinifera* (F6GV10) for Vitales, and *Populus trichocarpa* (A0A2K2BXT0) for Malpighiales. Only species with a single ZCCHC motif, whose CX_2_CX_4_HX_4_C consensus sequence is underlined in red, were selected for analysis. Asterisks and dots indicate identical (also shaded in black) and conserved residues, respectively.

**Supplementary Figure S7.** Predicted localizations of Arabidopsis CXIP4 and human SREK1P1 proteins. (A, B) Predicted (A) subcellular and (B) suborganellar localizations of CXIP4 and SREK1P1, using the MULocDeep web server (https://mu-loc.org/).

**Supplementary Figure S8.** Defects in 23S rRNA maturation in *cxip4-2* plants. (A) Schematic representation of the chloroplast *rrn* operon, indicating the lengths of the polycistronic primary transcript (purple), various processing intermediates (blue), and the 4.5S, 5S, 16S, and 23Sa/b/c mature rRNAs (red). Black and gray boxes represent chloroplast rRNA and tRNA genes, respectively, while lines indicate the intergenic sequences. The positions of the internal cleavage sites or hidden breaks (HB) in the 23S rRNA are shown with green arrowheads above the *rrn23* gene. The diagram was modified from Li *et al*. (2021). (B) Agilent 2100 bioanalyzer electropherogram profiles of total RNA extracted from Col-0 and *cxip4-2* plants collected 14 das. Peaks corresponding to chloroplast 4.5S/5S, 16S, and 23Sa/b/c rRNAs and cytoplasmic 18S and 25S rRNAs are labeled. Asterisks in the electropherogram profile of *cxip4-2* highlight the accumulation of incompletely processed 23S rRNAs of 2.4 (*) and 2.9 (**) kb.

**Supplementary Table S1.** Summary of protein-coding genes assigned to the PTHR31437 family according to the PANTHER database.

**Supplementary Table S2.** Summary of high-confidence physical protein-protein interactions for human SREK1IP1, identified from high-throughput experiments according to the HitPredict database.

**Supplementary Table S3.** Oligonucleotides used in this work.

**Supplementary Data Set S1.** Deregulated genes in *cxip4-2* plants.

**Supplementary Data Set S2.** Alternative splicing events in *cxip4-2* plants.

## Notes

### Competing Interest Statement

The authors have declared no competing interest.

### Summary of Updates

Supplementary Figures S1 to S8 added. Supplementary Tables S1 to S3 added. Supplementary Data Sets S1 and S2 added.

