## Supplementary Figures S1 to S8 and Supplementary Table S3 for "CXIP4 depletion causes early lethality and pre-mRNA missplicing in Arabidopsis"

The author responsible for distribution of materials integral to the findings presented in this article in accordance with the policy described in the Instructions for Authors (https://academic.oup.com/plphys/pages/General-Instructions) is María Rosa Ponce

### Supplementary Figures and Tables

Supplementary Material included in this file:

- Supplementary Figures S1 to S8
- Supplementary Table S3

Supplementary Material not included in this file:

- Supplementary Tables S1 and S2
- Supplementary Data Sets S1 and S2

CXIP4
MPATAGRVRMPANNRVHSSAALQTHGIWQSAIGYDPYAPTSKEEPKTTQQKTEDPENSYASFQGLLALARITGSNNDEARGS
CKKCGRVGHLTFQCRNFLSTKEDKEKDPGAIEAAVLSGLEKIRRGVGKGEVEEVSSEEEEESESSDSDVDSEMERIIAERFG
KKKGGSSVKKTSSVRKKKKRVSDESDSDSDSGDRKRRRRSMKKRSSHKRRSLSESEDEEEGRSKRRKERRGRKRDEDDSDES
EDEDDRRVKRKSRKEKRRRRSRRNHSDDSDSESSEDDRRQKRRNKVAASSDSEANVSGDDVSRVGRGSSKRSEKKSRKRHHR
KERE

CXIP4-1
MPATAGRVRMPANNRVHSSAALQTHGIWQSAIGYDPYAPTSKEEPKTTQQKTEDPENSYASFQGLLALARITGSNNDEARGS
CKKCGRVGHLTFQCRNFLSTKEDKEKDPGAIEAAVLSGLEKIRRGVGKGEVEEVSSEEEEESESSDSDVDSEMERIIAERFG
KKKGGSSVKKTSSVRKKKKRVSDESDSDSDSGDRKRRRRSMKKRSSHKRRSLSESEDEEEGRSKRRKERRGRKRDEDDSDES
EDEDDRRVKRKSRKEKRRRRSRRNHRIYSIVNGFMSGKSTWISNEYDGQYGEKERVITNFFSIQKCRCPQRYYKMKVHFDKT
TNYDPSYL

CXIP4-2
MPATAGRVRMPANNRVHSSAALQTHGIWQSAIGYDPYAPTSKEEPKTTQQKTEDPENSYASFQGLLALARITGSNNDEARGS
CKKCGRVGHLTFQCRNFLSTKEDKEKDPGAIEAAVLSGLEKIRRGVGKGEVEEVSSEEEEESESSDSDVDSEMERIIAERFG
KKKGGSSVKKTSSVRKKKKRVSDESDSDSDSGDRKRRRRSMKKRSSHKRRSLSESEDEEEGRSKRRKERRGRKMKP

**Supplementary Figure S1.** Localization of the ZCCHC motif, R-rich region, and nuclear and nucleolar localization signals in the predicted wild-type CXIP4 and mutant CXIP4-1 and CXIP4-2 proteins. The ZCCHC motif is highlighted in green, with the conserved cysteine and histidine residues ($CX_2CX_4HX_4C$) shown in red letters. The R-rich region, as annotated in UniProtKB (https://www.uniprot.org/uniprotkb), is underlined. Nuclear localization signals (NLSs) are denoted with red letters, and nucleolar localization signals (NoLSs) are highlighted in yellow. The 65 aa at the C-terminus of CXIP4-1 (blue letters) replace the 61 aa of the corresponding region of CXIP4. CXIP4-2 contains 240 aa, with only the last three absent from CXIP4 (blue letters).

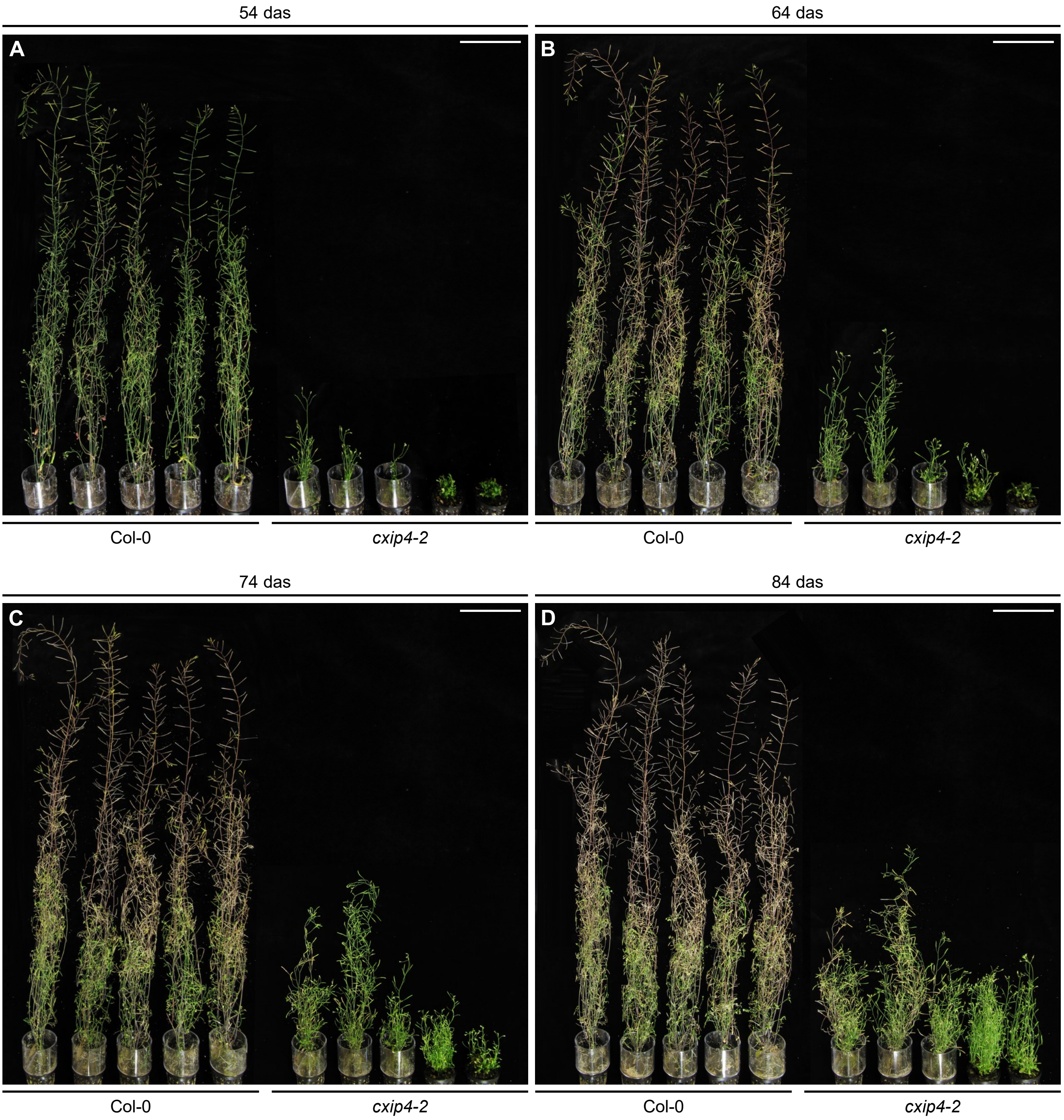


**Supplementary Figure S2.** Reproductive development of Col-0 and *cxip4-2* plants over time. Photographs were taken (A) 54, (B) 64, (C) 74, and (D) 84 das. Scale bars: 10 cm.

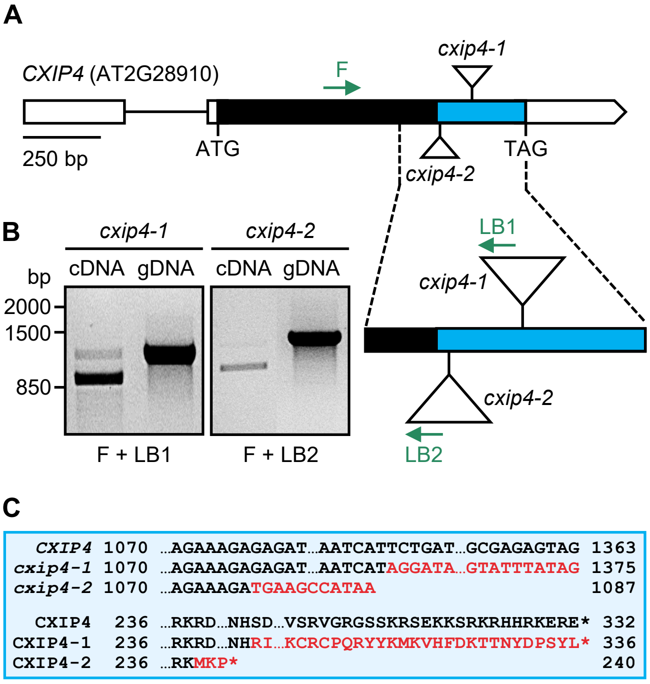


**Supplementary Figure S3.** Molecular effects of the T-DNA insertions in the *CXIP4* transcript. (A) Schematic representation of the *CXIP4* gene, as described in the Figure 1 legend, showing the positions of the primers used to analyze the effects of the T-DNA insertions (Supplementary Table S3). A segment of the gene is highlighted in blue and enlarged to better visualize such effects. Primers were not drawn to scale and are labeled as F (AT2G28910-F), LB1 (o8409), and LB2 (LBb1.3). (B) Agarose gel showing the PCR products obtained using cDNA and genomic DNA (gDNA) from *cxip4-1* and *cxip4-2* plants as templates and the F + LB1 and F + LB2 primer pairs. (C) Sequences of the chimeric *cxip4-1* and *cxip4-2* cDNAs and their predicted translation products CXIP4-1 and CXIP4-2. Nucleotide sequences corresponding to T-DNAs and the aa they are predicted to encode are highlighted in red. Asterisks indicate translation stop codons.

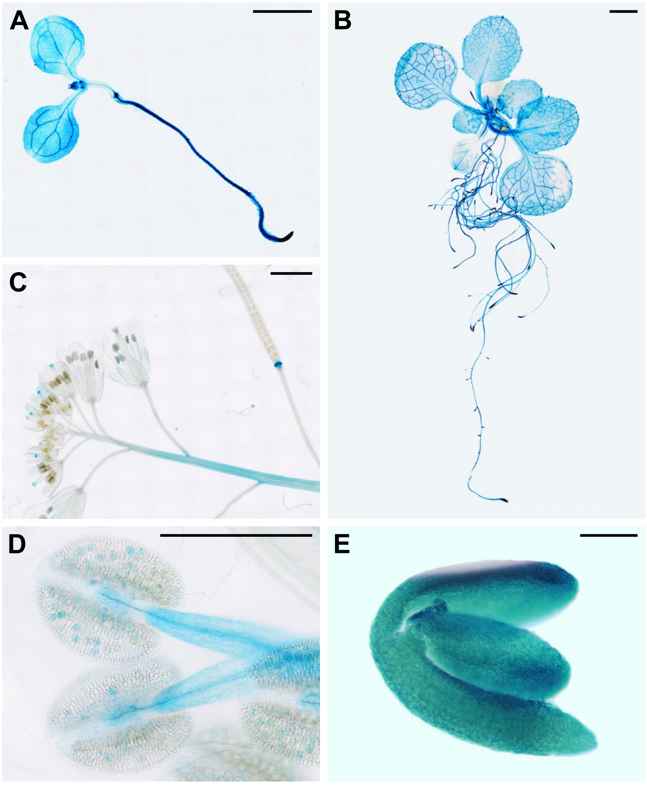


**Supplementary Figure S4.** Spatial expression of *CXIP4* during Arabidopsis development. GUS activity in Col-0 $\textit{CXIP4}_{\textit{proII}}\textit{:GUS}$ transgenic plants carrying the short region of the *CXIP4* promoter during (A, B) vegetative and (C-E) reproductive development. GUS staining was performed using (A, B) whole plants collected (A) 7 and (B) 14 das; (C, D) inflorescences, siliques, and anthers from 50 das-old plants; and (E) embryos dissected from green mature seeds of 52 das-old plants. Scale bars: (A-D) 2 mm, and (E) 100 µm. Equivalent GUS staining results were obtained from Col-0 $\textit{CXIP4}_{\textit{proI}}\textit{:GUS}$ plants.

```
C. reinhardtii     1 MATRGIAQLKAQYNRVRTSTALQGNSMWNNVIGIENEQKG-TNSAIEAALHAPSYGGGHK
A. thaliana        1 MPATAGRVRMPANNRVHSSAALQTHGIWQSAIGYDPYAP--TSKEEPKTTQQ-------K
P. patens          1 MPATAGRVRMPANNRVHSSAALQTHGIWQSAIGYDPYAPDKVQRQDDRSVPDPAAGGGAS
D. melanogaster      ------------------------------------------------------------
H. sapiens           ------------------------------------------------------------
Consensus

C. reinhardtii    60 AQRMERAQALIEQAGYQPKATDLQGLLALAKSQGATGNATRGACKICGGLGHLTKKCKNG
A. thaliana       52 TEDPENSYA------------SFQGLLALARITGSNNDEARGSCKKCGRVGHLTFQCRNF
P. patens         61 AADEQNAYD------------SFQGLLALARLTGSNADEARGSCQKCGRVGHLTFQCRNF
D. melanogaster    1 ---------------------------MNFPLASVNKDTLRAACKKCGYAGHLTYQCRNF
H. sapiens         1 -----------------------------MAVPGCNKDSVRAGCKKCGYPGHLTFECRNF
Consensus                                             . . :  *..*: **  **** :*.*

C. reinhardtii   120 VSGHTGDIGD-------LDAAAAVASMRALLPDPDEVSSLGSSDLDGSDLSDSDGDGGEK
A. thaliana      100 LSTKEDKEKDPGAIEAAVLSGLEKIRRGV---GKGEVEEVSSEEEEESESSDSDVDSEME
P. patens        109 LTAKEEAAAA-------AVSALERERNSAKFEGAGNLLA-ESSSSEESEISDSDEDSEME
D. melanogaster   34 LKVDPNKE---------ILLDVESTSSDSELDYLTPLTELRAQELKSGAEVPPPTEPAVL
H. sapiens        32 LRVDPKRD---------IVLDVSSTSSE-----------------------DSDEENEEL
Consensus            :                                                   .  :

C. reinhardtii   173 RKRKHSSSKKE-------------------------------------------------
A. thaliana      157 RIIAERFGKKKGGSSV--------------------------------------------
P. patens        161 RALAKLGRSKKGSSSVPDRKLSESKHSKSSSRKSKKKRHVSSSEFSDESSDGSRDRKRHR
D. melanogaster   85 PAAGKRDRSK--------------------------------------------------
H. sapiens        60 NKLQALQEKR--------------------------------------------------
Consensus                    ..

C. reinhardtii   184 -------KKEKDKKRKKEKSSKKGKKE---------------RKEKKDKREKRDKKRRHE
A. thaliana      173 -------KKTSSVRKKKKRVSDESDSD----------------------SDSGDRKRRRR
P. patens        221 KHRHSSKKKSSSRRHRSRRTKDDSDTDPSSDVEPESRHRHSKGRHLKDRKDRRSEKKRDA
D. melanogaster   95 -------DKSRDRLKAKKRERE---------------------------REREKVKEKEK
H. sapiens        70 -------------------------------------------------INEEEEKKKEK
Consensus                                                              :  . * .

C. reinhardtii   222 EDDREGSGKRARDESDSSSDSGSDSDGDRRREKRR-------RSSRERDSKEREQRGGRE
A. thaliana      204 SMKKRSSHKRRSLSESEDEEEG--------RSKRR-------KERRGRKRDEDDSDESED
P. patens        281 DYSDHDDRKKRKVEKSRDVEEGEIEDQRERVAKRRHDVESDIEDRRGKRSEKRRHDVDSD
D. melanogaster  121 EKGSRSKDKKRSHSKSSHKLAEKSKDKKSVRHKKH-------GKKRSRKHKKTTNTNNSS
H. sapiens        81 SKEKIKLKKKRKRSYSSSSTEEDTSKQKKQKYQKK-------EKKKEKKSKSK-------
Consensus            .       *.   . .                :..        . . . ..

C. reinhardtii   275 EDAREERRGEGHEDRRGEGHEDRRGEGREERR----------GEAREPEREREREHERER
A. thaliana      249 EDDRRVKRKSRKEKRR-----------------------------RRSRRNHSDDSDSES
P. patens        341 LEDQREKRPEKRGHDVDSDLDDRREIRSEKRRHDVDSDLDDRREIRSEKRRHEVDNDLED
D. melanogaster  174 SNSSSELAKTKTGKRPSSGSTNKKSKRRRSSSTSSSD-----SSSSSSSSSSSSSSTSSS
H. sapiens           ------------------------------------------------------------
Consensus

C. reinhardtii   325 EREHGRGGGRDEGEREYDREAWRRMERQRGEREGGAGGRNRDRDREERERGKEREQGRER
A. thaliana      280 SEDDRRQKRRNKVAAS-------------SDSEANVSGDDVSRVGRGSSKRSEKK-SRKR
P. patens        401 QRDKRSEKRRDVENSDFEDLRDKRVESTMSRMEDSEDGDERHRRKDRDERRSRRGGSRDR
D. melanogaster  229 SSSSSDDSSDDSSTSSDSSESDSSDNEESSTSESDEYGRKKKRQGKYKRKSTDTAMLRRK
H. sapiens       127 --------------------------------------KGKHHKKEKKKRKKEKHSSTPN
Consensus                                                      .      . .       .

C. reinhardtii   385 DRSHERDGSRERR--------------------
A. thaliana      326 HHRKERE--------------------------
P. patens        461 RHKERREYR------------------------
D. melanogaster  289 KTKAKRKRHRAGGSGAPTAGSSYLSSLSDSSTY
H. sapiens       149 SSEFSRK--------------------------
Consensus                 *.
```

**Supplementary Figure S5.** Sequence conservation among putative eukaryotic CXIP4 orthologs. Alignment of full-length putative CXIP4 orthologs from *Chlamydomonas reinhardtii* (UniProtKB accession number A0A2K3DE08), *Arabidopsis thaliana* (Q84Y18), *Physcomitrium patens* (A0A2K1K6W5), *Drosophila melanogaster* (Q9W3Z5), and *Homo sapiens* (Q8N9Q2). Asterisks and dots in the consensus line indicate identical (also shaded in black) and conserved residues, respectively. The consensus sequence of the ZCCHC motif is underlined in red.

```
H. vulgare          1 MPATAGRVRMPANNRVHSSAALQTHGIWQSAIGYDPYAPENNKQQPPSSSVSANAAAAAA
A. thaliana         1 MPATAGRVRMPANNRVHSSAALQTHGIWQSAIGYDPYAPTSKEEP---------------
C. annuum           1 MPATAGRVRMPANNRVHSSAALQTHGIWQSAIGYDPYAPSKDEDK---------------
V. vinifera         1 MPATAGRVRMPANNRVHSSAALQTHGIWQSAIGYDPYAPNKDDAK---------------
P. trichocarpa      1 MPATAGRVRMPANNRVHSSAALQTHGIWQSAIGYDPYAPNKDDSK---------------
Consensus             *************************************** ...

H. vulgare         61 NAAAEAAAPPSSDPGASSENAYTSFQGLLALARVTGSNSDETRGACKKCGRVGHLTFQCR
A. thaliana        46 ----KTTQQKTEDP----ENSYASFQGLLALARITGSNNDEARGSCKKCGRVGHLTFQCR
C. annuum          46 ----KSSTQKASSA-ADPENAYASFQGLLALARITGSNADETRGACKRCGRVGHLTFQCR
V. vinifera        46 ----NSSQPKSSNSEPEAENAYASFQGLLALARITGSNADETRGACKKCGRVGHLTFQCR
P. trichocarpa     46 ----KSSQEKSSTAEPEGENAYASFQGLLALARITSSNADEARGACKRCGRVGHLAFQCR
Consensus                 :::   :. .    **:*:**********:*.** **:**:**.*******:****

H. vulgare        121 NFLSVKELAMDDDVQAGMLSAAQAKAKFDEITKKASGARDGDEEAS-DEEDEDEIDSDSS
A. thaliana        98 NFLSTKE---DKEKDPGAIEAA-VLSGLEKIRRGV-----GKGEVE-EVSSEEEEESESS
C. annuum         101 NFVSVK----DDNKDAEAIEAA-VLSGLEKIKGNGSK---MKGKAENEESSEEEEESESS
V. vinifera       102 NFLSIKD---DKEKDPDAIQAS-VLSGLDKLKGNVGKKVNGKGAVEDESESEEEDESESS
P. trichocarpa    102 NFVSVKD--GDKEKDAEAVQAA-VLSGLEKIKGN------GKRVVQGEESEDDEDESETS
Consensus             **:* *    *.: :.  :.*: . : ::::          .  .. : ..::* :*::*

H. vulgare        180 DSDIDPELEKIIAERERAKSRGGKR-------------SGEEDKKASRHRTNSRGRSKHR
A. thaliana       148 DSDVDSEMERIIAERFGKKKGGSSVKKTSSVRKKKKRVSDESDSDSDSGDRKRRRRSMKK
C. annuum         153 DSDYDSEMERAIAEKYGKK--VSRKLKSSSKKHKKKDFDDDDDDESDSGKRKKRGRSKRR
V. vinifera       158 DSDADSEIERIIAERYEKK--GSSKRRSSRK-------EVSDDDGSDSRERKKRGRSKRR
P. trichocarpa    153 DSEEDSEIERIIAERYGKKK-SSTLGKSSRKK------EDSDDDGSDSG-RKKRGRSKKR
Consensus             **: *.*:*. ***.   *   .               . ..*. :.    : * **  ..

H. vulgare        227 RS-----RKSDSEDDSEEERTKDKKKKRRKKHKSSDEDSVSDSD----RKRHRKSRKDRK
A. thaliana       208 RSSHKR-RSLSESEDEEEGRSKRRKERRGRKRDEDDSDESEDED---DRRVKRKSRKEKR
C. annuum         211 RSGKKRGHSDSEDDDEDKERRKRRREKRRRRDDSSDED--EDED---RRRRRRKSRKEKR
V. vinifera       209 SGKR---VSGDLDDDDDESRRKRRKEKRKKRNDSSDEEKSSDEDKEERRLRRRKSRKEKR
P. trichocarpa    205 RIKK---RGSSDSEDGDEDRRKRRKEKRRKKNESSDEE----DD---RQRRKRKSRKEKR
Consensus                       . .:* :: * * ..:.* .. ...*.:    .*    .  .*****:..

H. vulgare        278 RRRTHRRKDDSSDDDESGGEGRRHHRHHKRRHHRRDASGSDSDGSESPHER-KRSSKQKS
A. thaliana       264 RRRSRRNHSDDSDSESSEDDR-R----QKRRNKVAASSDSEANVSGDDVSRVGRGSSKRS
C. annuum         266 RRRSHR-HANSSAESSDESPP-R----HKRRSRRAAASASDSDANNSDDARVGRD-KKRS
V. vinifera       266 RRRSHR-YSDDSDSSGDSD---R----HKRKSRR-AASLSDSDISGSDDSRVGRG-TKRS
P. trichocarpa    255 RRRSHR-HSDDSDSDEHEHSGRR----HKQKSRK-ASSLSDPDASGSDDPRVGRD-TKRS
Consensus             ***:.*   :.* ..       *    :*.. .   :* *:.: . .   *  *. .:.*

H. vulgare        337 HKRSESHGLGKDERHCPQGASRSGEKSRG
A. thaliana       319 EKKSR---------------KRHHRKERE
C. annuum         319 EKRSR---------------KRHDDEE--
V. vinifera       316 EKRNR---------------KRHHKLDD-
P. trichocarpa    308 EKKIR---------------KRHHEDDE-
Consensus              *.                 .*    .
```

**Supplementary Figure S6.** Sequence conservation among CXIP4 orthologs in angiosperms. Alignment of full-length CXIP4 orthologs from different angiosperm lineages: *Hordeum vulgare* (UniProtKB accession number A0A287R5P6) representing Poales, *Arabidopsis thaliana* (Q84Y18) for Malvids, *Capsicum annuum* (A0A2G3ANM6) for Asterids, *Vitis vinifera* (F6GV10) for Vitales, and *Populus trichocarpa* (A0A2K2BXT0) for Malpighiales. Only species with a single ZCCHC motif, whose $CX_2CX_4HX_4C$ consensus sequence is underlined in red, were selected for analysis. Asterisks and dots indicate identical (also shaded in black) and conserved residues, respectively.

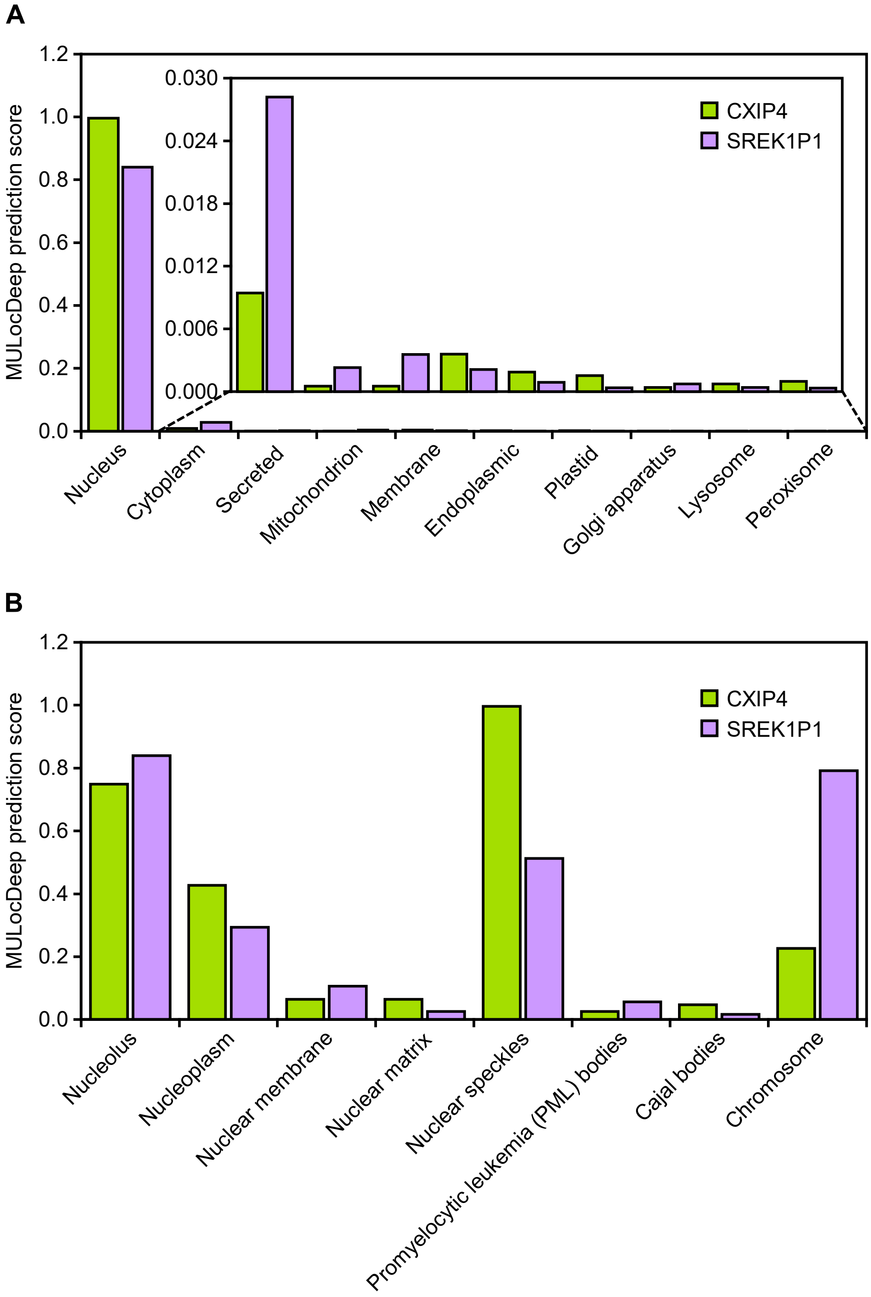


**Supplementary Figure S7.** Predicted localizations of Arabidopsis CXIP4 and human SREK1P1 proteins. (A, B) Predicted (A) subcellular and (B) suborganellar localizations of CXIP4 and SREK1P1, using the MULocDeep web server (https://mu-loc.org/).

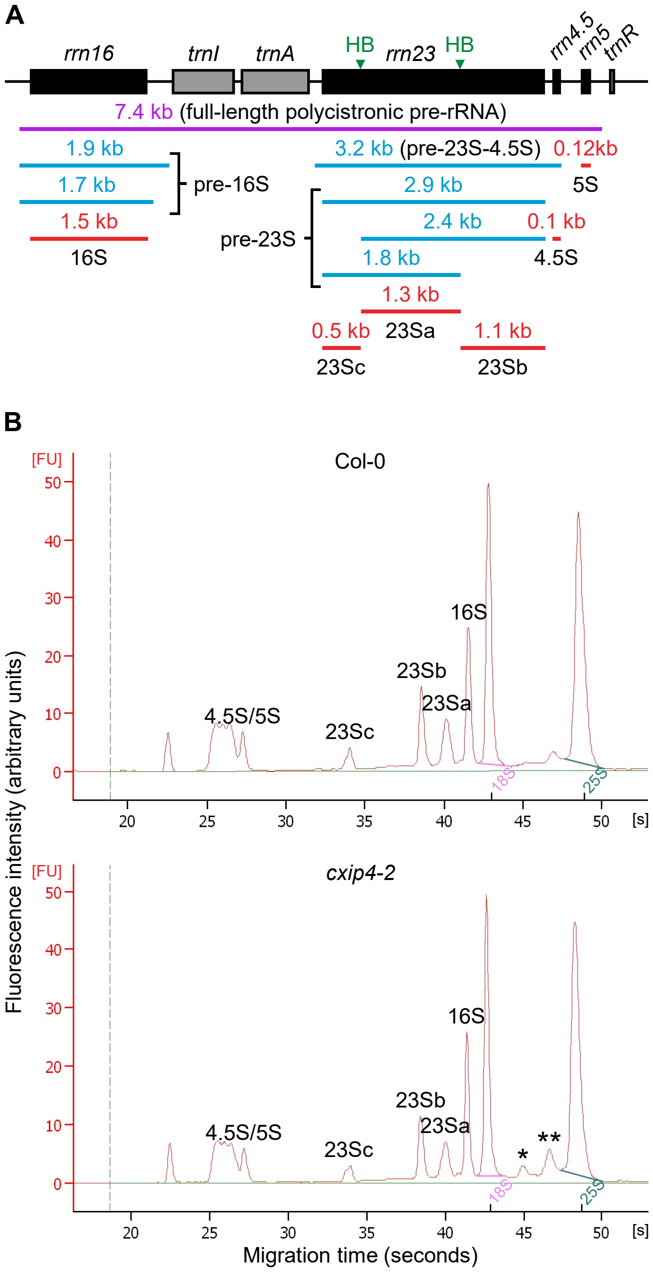


**Supplementary Figure S8.** Defects in 23S rRNA maturation in *cxip4-2* plants. (A) Schematic representation of the chloroplast *rrn* operon, indicating the lengths of the polycistronic primary transcript (purple), various processing intermediates (blue), and the 4.5S, 5S, 16S, and 23Sa/b/c mature rRNAs (red). Black and gray boxes represent chloroplast rRNA and tRNA genes, respectively, while lines indicate the intergenic sequences. The positions of the internal cleavage sites or hidden breaks (HB) in the 23S rRNA are shown with green arrowheads above the *rrn23* gene. The diagram was modified from Li *et al.* (2021). (B) Agilent 2100 bioanalyzer electropherogram profiles of total RNA extracted from Col-0 and *cxip4-2* plants collected 14 das. Peaks corresponding to chloroplast 4.5S/5S, 16S, and 23Sa/b/c rRNAs and cytoplasmic 18S and 25S rRNAs are labeled. Asterisks in the electropherogram profile of *cxip4-2* highlight the accumulation of incompletely processed 23S rRNAs of 2.4 (*) and 2.9 (**) kb.

**Supplementary Table S3.** Oligonucleotides used in this work

| Purpose | Names | Sequences (5′ → 3′) |
|---|---|---|
| Genotyping of *cxip4-1* and *cxip4-2* insertional alleles | AT2G28910-F[a] | GTCGGGGTTGGAGAAGATTAG |
| | AT2G28910-R[a] | TATGGTTCAACGGTGGAGAAG |
| | o8409[a,b] | ATATTGACCATCATACTCATTGC |
| | LBb1.3[a,c] | ATTTTGCCGATTTCGGAAC |
| Gateway cloning and verification of constructs | CXIP4proI-F[d] | *GGGGACAAGTTTGTACAAAAAAGCAGGCT*AAAGTTCCACATTGCCCTCTAC |
| | CXIP4proII-F[d] | *GGGGACAAGTTTGTACAAAAAAGCAGGCT*GAGTTTATTGAATTGTGGGGATG |
| | CXIP4proI/II-R[d] | *GGGGACCACTTTGTACAAGAAAGCTGGGTG*CTTTTCGAAACCCTAGAAAAAAC |
| | CXIP4proII:CXIP4:GFP-R[d] | *GGGGACCACTTTGTACAAGAAAGCTGGGTG*CTCTCGCTCTTTCCTGTGATG |
| | M13-F | GTTGTAAAACGACGGCCAGTG |
| | GUS-R | CACAAACGGTGATACGTACACT |
| | GFP-R | CTTGTACAGCTCGTCCATGC |

The oligonucleotides used as forward and reverse primers in PCR amplifications and/or Sanger sequencings are indicated as -F or -R, respectively. [a]The *CXIP4* wild-type allele was PCR amplified with the [a]AT2G28910-F/R primers, flanking the T-DNA insertions of *cxip4-1* and *cxip4-2*. The *cxip4-1* and *cxip4-2* insertional alleles were amplified using the AT2G28910-F + o8409 and AT2G28910-F + LBb1.3 primer pairs, respectively. [b,c]The sequences of these primers were taken from [b]https://www.gabi-kat.de/faq/vector-a-primer-info.html and [c]http://signal.salk.edu/tdnaprimers.2.html. [d]The *att*B sequences used for the Gateway cloning of the PCR products are shown in italics.
